# Coordinated protein modules define DNA damage responses to carboplatin at single cell resolution in human ovarian carcinoma models

**DOI:** 10.1101/2024.11.21.624591

**Authors:** Jacob S. Bedia, Ying-Wen Huang, Antonio Delgado Gonzalez, Veronica D. Gonzalez, Ionut-Gabriel Funingana, Zainab Rahil, Alyssa Mike, Alexis Lowber, Maria Vias, Alan Ashworth, James D. Brenton, Wendy J. Fantl

## Abstract

Tubo-ovarian high-grade serous carcinoma (HGSC) is the most lethal gynecological malignancy and frequently responds to platinum-based chemotherapy because of common genetic and somatic impairment of DNA damage repair (DDR) pathways. The mechanisms of clinical platinum resistance are diverse and poorly molecularly defined. Consequently, there are no biomarkers or medicines that improve patient outcomes. Herein we use single cell mass cytometry (CyTOF) to systematically evaluate the phosphorylation and abundance of proteins known to participate in the DNA damage response (DDR). Single cell analyses of highly characterized HGSC cell lines that phenocopy human patients show that cells with comparable levels of intranuclear platinum, a proxy for carboplatin uptake, undergo different cell fates. Unsupervised analyses revealed a continuum of DDR responses. Decompositional methods were used to identify eight distinct protein modules of carboplatin resistance and sensitivity at single cell resolution. CyTOF profiling of primary and secondary platinum-resistance patient models shows that a complex DDR sensitivity module is strongly associated with response, suggesting it as a potential tool to clinically characterize complex drug resistance phenotypes.

## Introduction

Cytotoxic chemotherapy remains a critically important treatment for cancer patients worldwide and is effective against rapidly dividing tumor cells in which the DNA damage response (DDR) is deregulated from cell cycle control ^1–3^. However, durable patient outcomes are rare, even in individuals with enhanced innate sensitivity. This is due to acquisition and selection of complex cellular traits that result in therapeutic resistance ^4,5^. The standard of care for women with tubo-ovarian high grade serous carcinoma (HGSC) remains carboplatin-based chemotherapy either following surgery or in a neoadjuvant setting ^6–11^. Most patients (∼ 60%) initially respond because of frequent impairment of DDR pathways but almost all will develop fatal platinum-resistant recurrent disease ^12,13^. The exact mechanisms behind clinical platinum resistance are not well understood and consequently, there are no effective medicines specifically designed to target carboplatin resistance.

Although multiple potential resistance mechanisms have been described, including genomic alterations, enhanced DNA repair capabilities and increased drug transporter activity, they have shown limited clinical impact ^9,14–16^. However, the importance of genetic reversion as a mechanism of resistance is strongly supported from clinical studies in patients with *BRCA1* and *BRCA2* mutations. These studies have demonstrated that tumors can acquire secondary somatic mutations that restore the function of homologous recombination (HR) proteins, thereby contributing to resistance both to carboplatin and poly (ADP-ribose) polymerase inhibitors (PARPi) ^17–21^. Although HGSC tumors are primarily driven by DNA structural variants, especially copy number alterations (CNA) ^22^ there is little evidence for new selection of commonly occurring oncogenic CNAs ^23^.

Additionally, methylation, gene expression integrated with other omics data reported by the Ovarian Cancer Genome Atlas (TCG) has highlighted the complexity of resistance without identifying any specific therapeutic targets ^24^. Single cell RNA expression studies show progressive stepwise adaptation toward resistance in response to PARPi treatment in ovarian cancer models. These trajectories are potentiated by epithelial to mesenchymal transition states and maintained by diverse reprogramming of metabolic and stress phenotypes ^25^. Together these studies suggest that HGSC tumors are susceptible to a broad evolutionary space to escape carboplatin and PARPi therapy.

Although the potential upstream genetic, epigenetic and transcriptomic effects are highly diverse, they necessarily must converge on protein function within the DDR network. Critically, the DDR is regulated by protein expression levels and phosphorylation states ^26–30^. Based on the reported heterogeneity of resistance phenotypes and possible DDR responses, we hypothesized that: i) single cell proteomic analysis is required to identify diverse DDR protein networks indicative of sensitivity or resistance to carboplatin and ii) comparable levels of carboplatin uptake by individual cells will result in different DDRs. To address our hypotheses we applied mass cytometry, known as CyTOF/<u>Cy</u>tometry by <u>T</u>ime-<u>O</u>f-<u>F</u>light to measure the DDR in HGSC tumor cells. CyTOF is a single cell proteomic technology that can measure up to 60 parameters per cell using a panel of antibodies tagged with heavy metal-chelating polymers ^31–33^. We assembled a CyTOF antibody panel to simultaneously measure the DDR, cell cycle phase and intracellular signaling with measurements of intracellular platinum in response to carboplatin alone or in combination with a PARPi. The atomic mass of platinum falls within the measurable mass range of CyTOF. Critically, our study used HGSC model systems that more closely represent primary tumors than previous models. Unsupervised analysis of millions of single cells discovered DDR protein modules recruited by cells in specific states after treatment.

## Results

### Validating a CyTOF antibody panel to measure the DDR and cell cycle

We designed a CyTOF antibody panel to measure the DNA damage response in individual cells throughout all cell cycle phases **(Fig. 1B, Supp. Table S1)**. Thirty five antibodies were validated for: DNA damage detection, non-homologous end joining (NHEJ) and homologous recombination (HR) repair processes, cell cycle phases, cell cycle checkpoint activation, cell cycle arrest and phosphorylated (activated) intracellular signaling pathways ^34^ **(Methods)**. Measurements also included a live-dead cell discriminator (either cisplatin or rhodium-103 [^103^Rh]) and a marker for apoptosis (cleaved (c)PARP) ^35,36^. All cell cycle phases were identified by gating with pRb(S807/811), IdU (demarcates cells in S-phase), cyclin E, cyclin B1, and pHH3(S28) ^37^ **(Supp. Fig. 1).** In a pilot study of HeLa cells treated with a variety of genotoxic agents overall, we observed expected DDRs that validated our panel. However, visualization of the single cell data with UMAPs revealed a previously unrecognized level of DDR complexity. Some DDRs were specific to cells within a particular cell cycle phase. In other cases, cells in the same cell cycle phase showed varying responses to the different agents used but also to a given specific agent **(Supp. Text and Supp. Figs. 2 and 3)**.

**Fig. 1:**
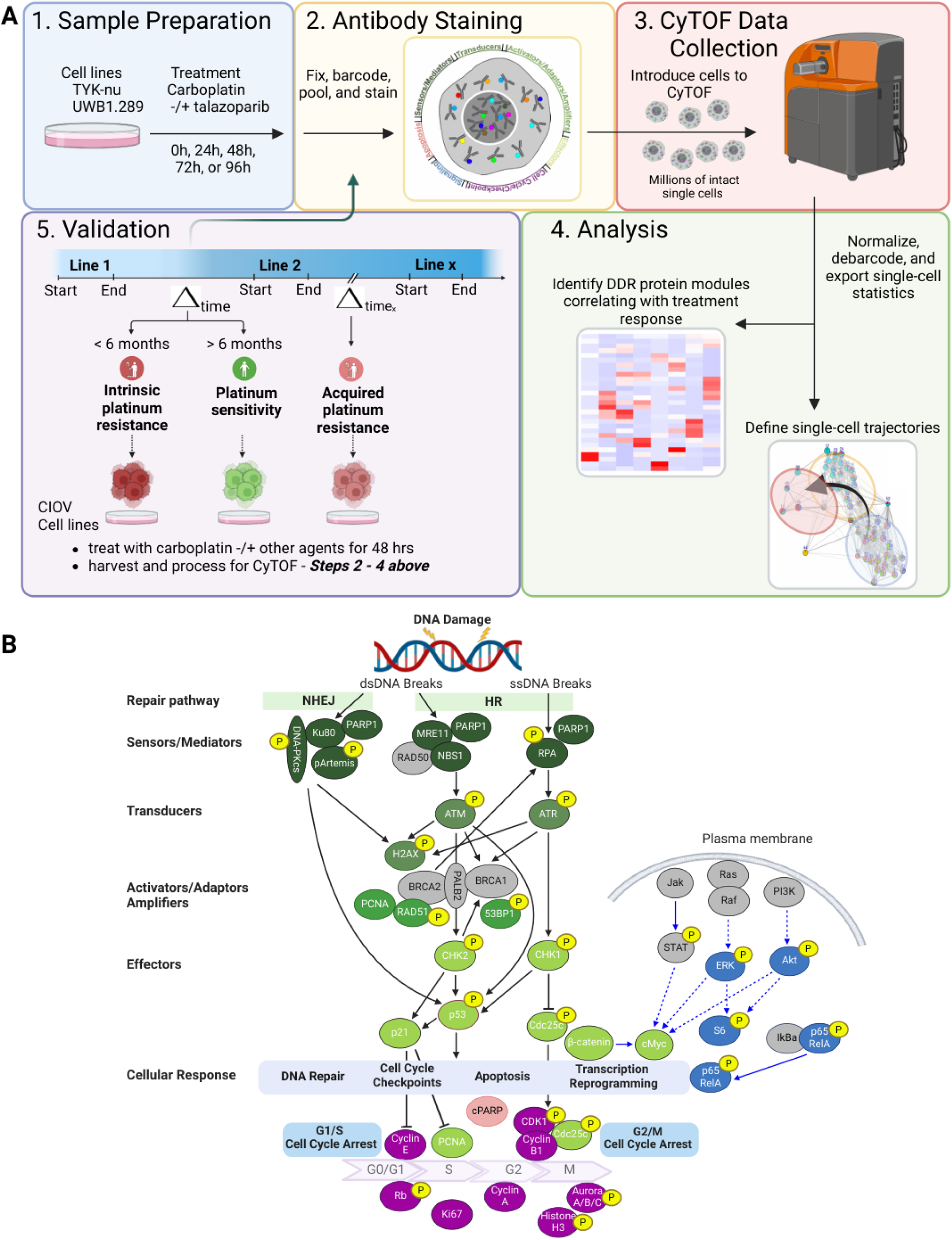
Characterization of the DNA damage response by CyTOF. **A.** Schema of experimental approach. Foundational experiments were performed using TYK-nu and UWB1.289 HGSC cell lines. Validation was performed using three spontaneously immortalized continuous HGSOC cell lines named CIOV1, CIOV2, and CIOV3. **B.** Pathway map showing CyTOF antibody panel designed to measure proteins participating in the carboplatin DNA damage response. Proteins marked in grey were not measured because no suitable antibodies were available. “β-catenin” is non-phospho-β-catenin **(Supp. Table 1** shows positive and negative controls for antibody validation).

### Modelling carboplatin resistance in clinically relevant HGSC cell lines

In a previous CyTOF study we characterized HGSC tumor cell suspensions disaggregated from freshly resected advanced-stage tumors, along with 13 molecularly characterized HGSC cell lines ^38–40^. Our data revealed that the HGSC cell lines reflected specific tumor cell phenotypes identified from the primary HGSC tumors. Notably, the TYK-nu cell line phenocopied poor-prognosis vimentin-expressing cells comprised of both carboplatin-resistant and responsive subpopulations ^38,39^. We therefore selected the TYK-nu cell line as well as the UWB.289-BRCA1-null cell line derived from a highly chemo-resistant HGSC tumor and its BRCA1-expressing counterpart to study the DDR response to carboplatin over time. We validated our results in three spontaneously immortalized HGSC cell lines (Cambridge Institute Ovarian (CIOV)1, CIOV2 and CIOV3) that retained the key characteristics of their original tumors and represented distinct states of platinum resistance ^41^ **(Fig. 1A)**.

The TYK-nu cell line was generated from a xenograft of a primary patient derived ovarian tumor and closely phenocopies the poor prognosis HGSC cell populations identified in newly diagnosed primary tumors ^39, 42^ **(Supp. Table 2)**. Both clinical and preclinical studies showed that combining PARP inhibitors with carboplatin-based chemotherapy significantly improved progression-free survival and efficacy ^43–45^. Specifically, preclinical studies demonstrated that carboplatin’s effectiveness was increased by inhibiting PARP’s enzyme activity and enhancing its DNA-trapping ability ^44,45^. We therefore designed CyTOF experiments to analyze DDRs in TYK-nu cells treated with carboplatin alone or in combination with PARPi over time. Since the DDR may be strongly altered by PARP DNA trapping activity, we chose to use talazoparib, as one of the most potent PARP trapping agents ^46–48^. Cells were exposed to either carboplatin alone (8μM), carboplatin (8μM) plus talazoparib (100nM), or talazoparib alone (100nM) for 24, 48, 72, and 96h ^9,11^. Optimal drug concentrations were selected from dose response curves **(Supp.** Fig. 4 and **Methods**). At each timepoint cells were incubated with ^103^Rh (a live-dead marker), barcoded, combined, stained with the antibody panel and processed for CyTOF **(Methods)** ^34^. Gating for live cells (negative for ^103^Rh), generated a CyTOF dataset of 721,579 cells for downstream analysis **(Supp. Table 3)**.

### Cell cycle responses to carboplatin induce S-phase

Each cell cycle phase was manually gated from the viable (^103^Rh-c-PARP-) TYK-nu cell population **(Fig. 2A and Methods)**. Under all conditions, the proportion of cells in G1 was < 1%, due to the abrogated G1 checkpoint caused by *TP53* mutation ^49^. All three treatments promoted a dramatic and maximal increase in the proportion of cells that arrested in S-phase, ∼80% at 24 and 48hr, consistent with a major DDR **(Supp. Table 3)**. At 72hr and 96hr, a proportion of cells had transitioned into G2, but a significant proportion remained in S-phase, which was most marked after carboplatin mono-treatment. Cells spent minimal time in M-phase. Although we could not determine whether cells in G0 had undergone therapy-induced senescence or quiescence, both states have been associated with drug resistant phenotypes ^50^.

**Fig. 2.**
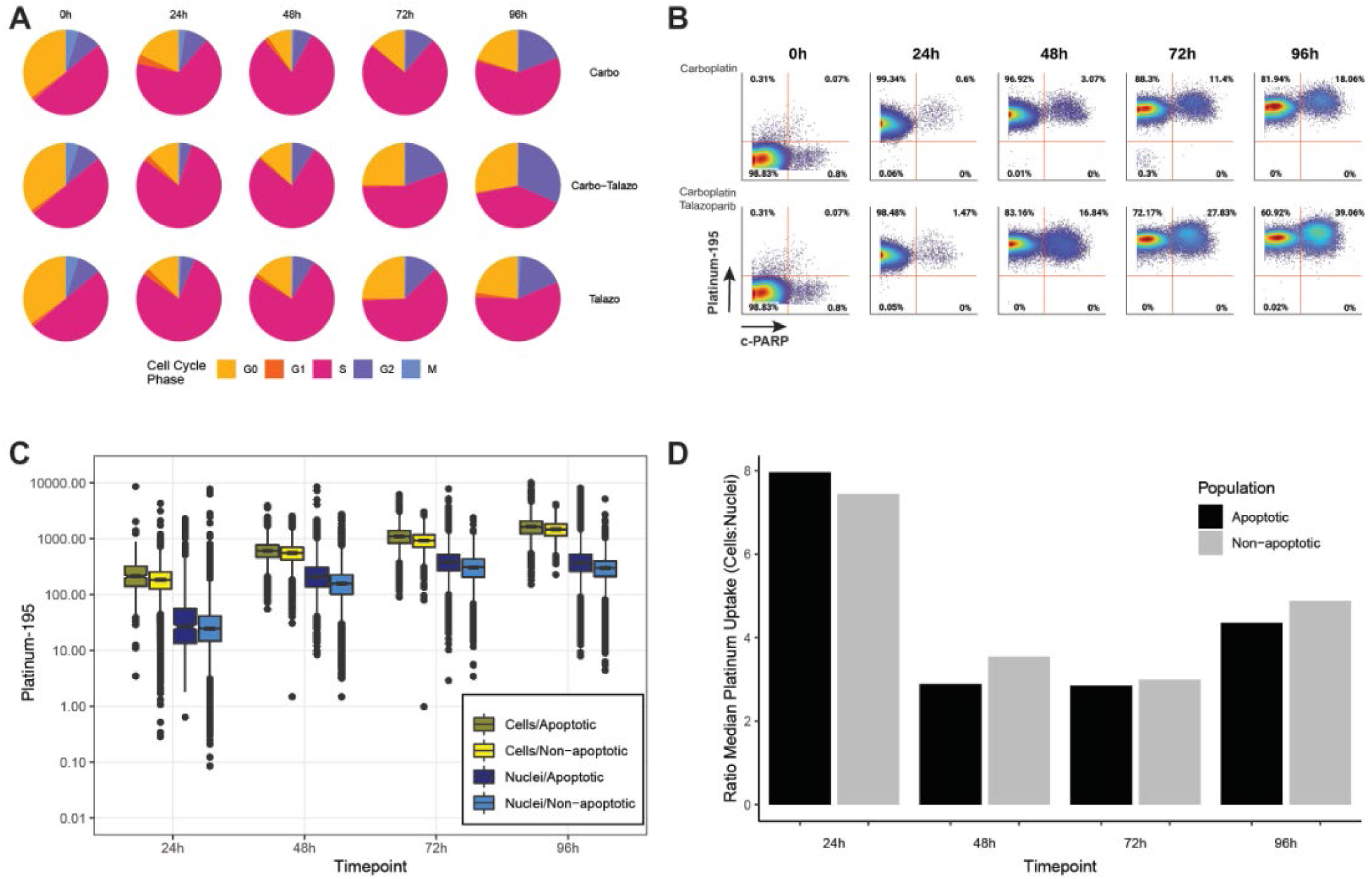
Characterization of responses to carboplatin in the TYK-nu cell line. TYK-nu cells were treated with carboplatin, talazoparib, or both drugs and processed for CyTOF analysis at the indicated times. **A.** Pie charts depict cell frequency distributions across cell cycle phases. **B.** Biaxial plots displaying ^195^Pt levels, which indicate carboplatin uptake, plotted against c-PARP levels to differentiate apoptotic from non-apoptotic cells to treatments over time. **C.** Box and whisker plot showing platinum uptake in single intact cells and single isolated nuclei over time. CyTOF enabled characterization of apoptotic populations at early times of drug treatments when cell frequencies were low (∼200 to 1000 cells). Boxes are colored by cell fate (apoptotic or non-apoptotic) and compartment (whole cell (yellow) or nucleus (blue)). Notches are calculated to give a 95% confidence interval comparing median values for ^195^Pt uptake. **D.** Fold change in median platinum levels comparing whole cells to nuclei for each timepoint and population.

### Quantifying non-apoptotic and apoptotic cell populations in response to carboplatin

CyTOF provides a unique opportunity to quantitatively measure intracellular levels of platinum (^195^Pt) ^51,52^. ^195^Pt measurements can be used as a surrogate for intracellular carboplatin levels (Methods). ^103^Rh-cells were manually gated with c-PARP to enumerate apoptotic and non-apoptotic cell populations after carboplatin (C) and carboplatin + talazoparib (C + T) treatments. Biaxial plots revealed that while the frequency of apoptotic cells increased over exposure time (18% for carboplatin alone, 39% for carboplatin plus talazoparib at 96h), a large population of cells remained non-apoptotic **(Fig. 2B)**. These data are consistent with previous studies showing that TYK-nu cells are comprised of cell populations with differing carboplatin sensitivities. They also show that talazoparib potentiates carboplatin activity by accumulating DNA damage through the inhibition of PARP-mediated DNA repair ^53–55^.

### Quantifying nuclear uptake of carboplatin

The 2D biaxial plots indicated that a proportion of apoptotic and non-apoptotic populations had comparable levels of carboplatin **(Fig. 2B)**. Box and whisker of individual cell concentrations for ^195^Pt confirmed increased carboplatin uptake over time but also revealed a previously unrecognized variability in intracellular ^195^Pt levels **(Fig. 2C)**. Although median uptake levels were greater for apoptotic cells, a large proportion of non-apoptotic cells within the interquartile range had the same level of carboplatin (69–89%) at each time point **(Fig. 2C)**. Additionally, some cells survived despite having extremely high levels of ^195^Pt uptake.

Measurements of cellular carboplatin include adduct formation between proteins and DNA ^56^. However, the carboplatin-mediated DDR necessarily depends on nuclear uptake and the formation of stable DNA adducts. We therefore developed a new CyTOF protocol to measure carboplatin levels in individual intact nuclei isolated from TYK-nu cells **(Supp.** Fig. 5 and **Methods)**. Nuclear uptake of carboplatin increased over time but at lower levels compared with cellular uptake **(Fig. 2C)**. The fold change of carboplatin uptake between cells and nuclei was equivalent for apoptotic and non-apoptotic cells at each time point, showing that alterations in nuclear import/export of carboplatin were not the main determinants of apoptotic cell fate **(Fig. 2D)**. As seen for total cellular uptake, the median carboplatin uptake was greater in the nuclei of apoptotic versus non-apoptotic cells. However, we also noted that significant numbers of nuclei from apoptotic and non-apoptotic cells had the same level of ^195^Pt (65–100%) at each timepoint. The data demonstrate that drug influx and efflux pumps play only a partial role in resistance.

### Mapping the DDR trajectory in single cells

These pharmacodynamic and pharmacokinetic data led us to hypothesize that cell populations with similar carboplatin levels, but different fates must have distinct DDRs. We therefore generated a pipeline to analyze DDR dynamics temporally and in an unbiased manner independent of treatment, cell cycle and cell fate **(Fig.3)**. We first computed a UMAP embedding of the TYK-nu single cell data from all timepoints and treatments using 29 DDR, cell cycle checkpoint and intracellular signaling proteins **(Fig. 4A, Supp.** Fig. 6). The UMAP embedding revealed a predominant continuum of cells with only three discernible cell populations. In contrast to UMAPs generated with phenotypic markers where discrete cell populations are easily visualized (e., g T cells, B cells), this UMAP was generated exclusively with intracellular functional markers (phospho-states, protein levels) revealing a continuum of many subtle and different DDR functional states. To identify cell subpopulations within the UMAP, we applied Leiden clustering to group cells into small neighborhoods based on their DDRs **(Fig. 4A)**. These clusters were visualized with different colors and each cluster’s centroid was labelled on the UMAP ^57^. Next, we used partition-based graph abstraction (PAGA) to map the relationships between these Leiden clusters. In the PAGA plot, each node represents a Leiden cluster, and the edges between nodes indicate the degree of connectivity between clusters. ^58^. To align the PAGA graphs with the UMAP, the PAGA nodes were positioned at the cluster centroids **(Fig. 4A-E)**. Each cluster in the PAGA graph was depicted by a pie chart showing the proportions of cells from different conditions.

**Fig. 3.**
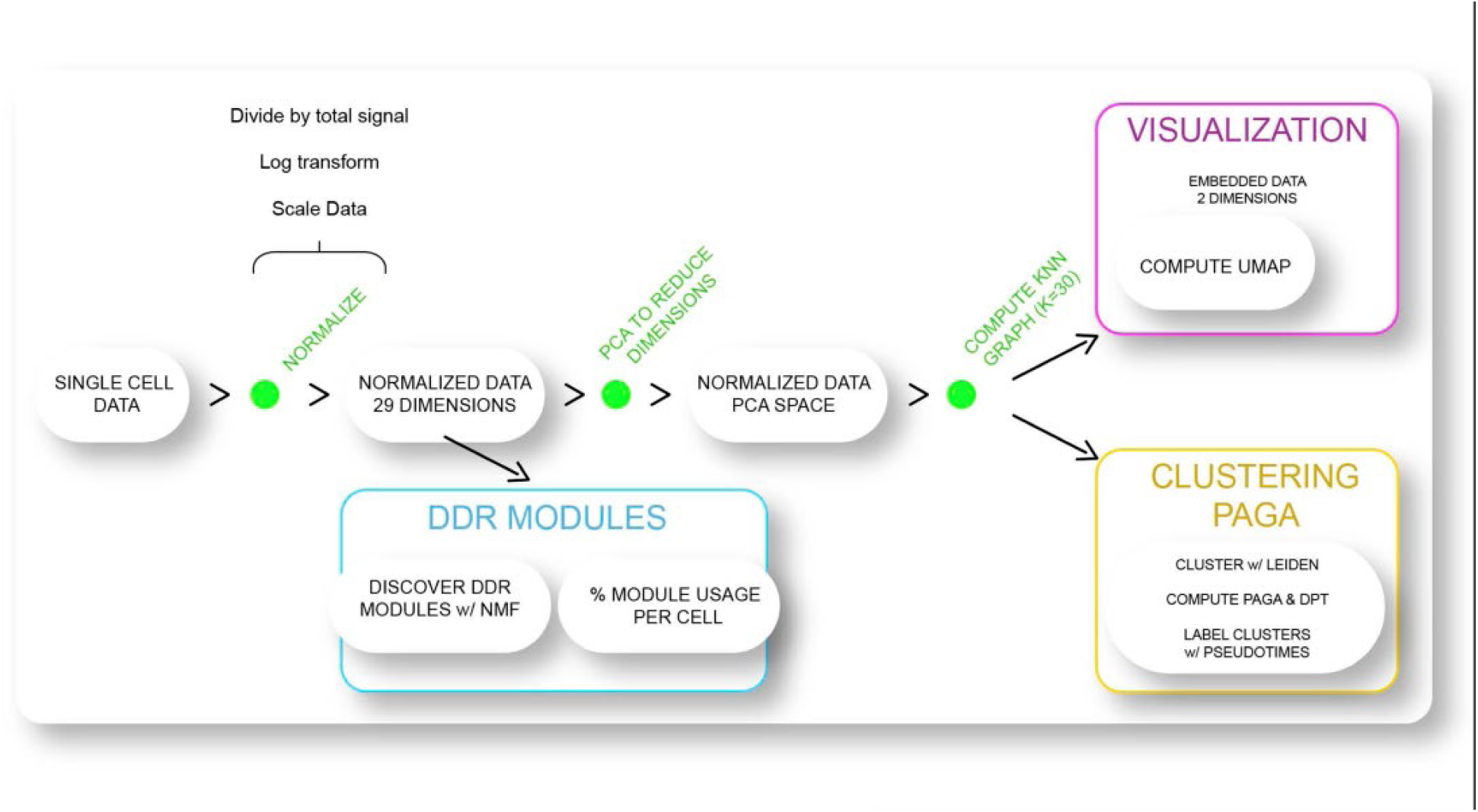
**Schema depicting unsupervised data analysis approach.**

**Fig. 4.**
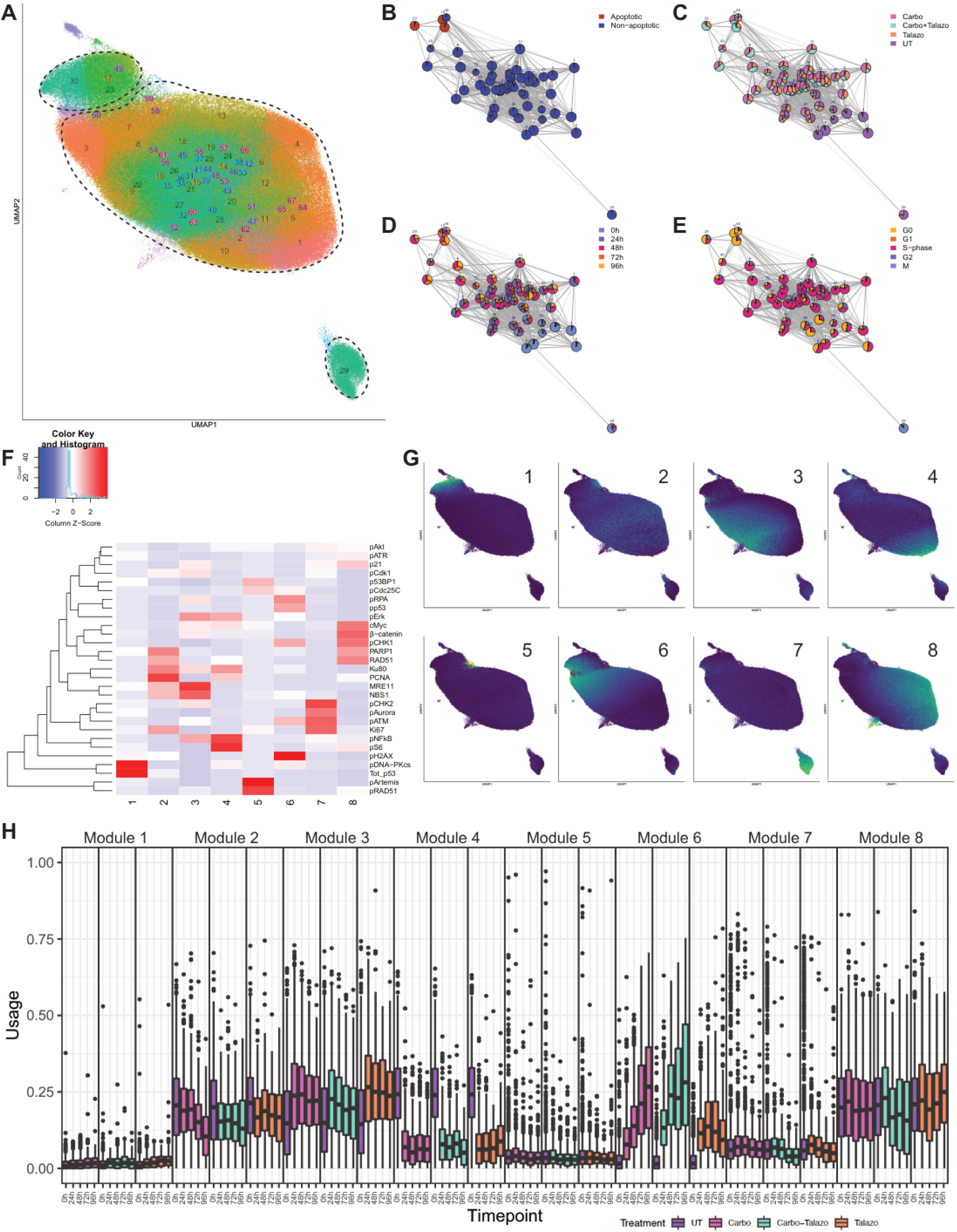
Identification of DDR protein modules in TYK-nu cells. **A**. UMAP embedding generated with 29 DDR proteins, of single cell data from all timepoints for 721,579 TYK-nu cells. Leiden cell clusters are overlaid on the UMAP and colored. **B–E.** Partition-based graph abstraction plots show connectivity between Leiden clusters. Plots are colored for cell fate, treatment, time, and cell cycle. Clusters are colored with a pie-chart showing proportion of cells with different DDR features. **F and G.** DDR protein modules discovered by non-negative matrix factorization (NMF). The matrix of expression levels for 29 DDR proteins in 721,579 cells was decomposed into two matrices. One discovered the most frequenctly co-occurring proteins in eight (number user selected) DDR modules. The contribution of each protein within a specific module is given by its z-score and this matrix is depicted on a hierarchically clustered heatmap. The second matrix describes the activity of each module in an individual cell and is overlaid on the UMAP. **H.** Box and whisker plots depict the activity of each module activity over time.

PAGA revealed that the mass of cells in the upper left of the UMAP were apoptotic with a DDR signature that was distinct from non-apoptotic cells **(Fig. 4A)**. Most untreated cells were in the lower right of the UMAP **(Fig. 4C)**. Upon treatment, as the effect of the agents increased, cells had a trajectory toward the apoptotic cell state. For example, when compared to cells treated with T alone, cells treated with C + T located further away from the untreated DDR and closer to the apoptotic state **(Fig. 4C)**. Mapping the PAGA graphs over time revealed a rough temporal progression. However, timepoints were not discrete. For example, some cells that had been treated with C + T for 96hr had a comparable DDR with untreated or treated cells at early time points. By contrast, some cells treated for 24hr mapped closely to cells treated for 96hr and in proximity to apoptotic cells **(Fig. 4D)**. These results demonstrated that cells are progressing and responding to treatment at different rates. Cell cycle analysis showed that most cells were in G0 or S-phase. Nevertheless, there was a clear signature of cells in M-phase on the lower righthand corner of the PAGA graph **(Fig. 4E)**.

### Non-negative matrix factorization (NMF) to discover modules of co-occurring DDR proteins

The UMAP/PAGA analysis revealed clusters of cells with different DDRs. To determine if specific protein modules influenced the positioning of cells along the PAGA trajectories, we applied non-negative matrix factorization (NMF) **(Fig. 4F)**. This algorithm simultaneously learns a set of co-occurring proteins within DDR modules and then computes the activity of each module in each cell ^59^. NMF computed eight DDR modules and the contribution of each protein to each module was then visualized with a heatmap **(Fig. 4F)**. Several proteins occupied more than one module; for example, PARP1 was part of Modules 2 and 8, and PCNA was part of Modules 2 and 4 **(Fig. 4F)**. To determine whether cells recruit different modules over time, these modules were overlaid on the PAGA graphs **(Fig. 4G)**. Rather than being uniformly distributed throughout the PAGA graphs, the overlays showed that most modules were confined to discrete populations of cells over the time-course. Four modules with high activity mapped mostly to localized regions of the PAGA graphs: apoptotic cells to Module 1, UT cells (endogenous DDR) to Module 4 to, cells treated for longer times to Module 6 to late timepoints and mitotic cells to Module 7.

### Quantifying DDR usage over time in single cells

To further characterize module usage shown in the overlays we generated box and whisker plots **(Fig. 4H)**. The plots showed that module activity changed over time at different rates. Modules 2 and 8 were active in both endogenous and exogenous DDRs at early timepoints and Module 4 was most active in untreated cells. Module 3 peaked early after treatment with MRE11 and NBS1 consistent with their role as early sensors of DNA damage. In contrast, Module 6 demonstrated the greatest change over time, bridging pre-apoptotic and apoptotic cells (**Fig. 4G, H**). Furthermore, Module 6 usage was largely driven by carboplatin and not by talazoparib in this study. Module 6 was comprised of pH2AX, pATM, pCHK1, pp53 and pRPA, all proteins playing critical roles in HR DNA double strand repair ^60^. The recruitment of Module 6 at later timepoints suggested it could be a pharmacodynamic marker of responsiveness to carboplatin-induced damage. The key finding from the UMAP/PAGA/NMF analysis was that while most non-apoptotic cells mapped together densely on the UMAP, functionally they could be distinguished by eight distinct DDR protein modules.

### Responses of UWB1.289 BRCA1- and BRCA1+ cell lines to carboplatin

To determine whether the carboplatin-mediated DDR modules identified in the TYK-nu cell line are conserved in other HGSC cells, we performed a carboplatin treatment time course and DDR-CyTOF analysis on the UWB1.289 BRCA1-/BRCA+ isogenic cell line pair **(Supp. Table 1)** ^61^. The UWB1.289 BRCA1- cell line was derived from an HGSC tumor that recurred after primary debulking surgery and treatment with carboplatin/paclitaxel, paclitaxel, topotecan, and gemcitabine with doxorubicin. It harbors a clinically deleterious allele of *BRCA1* and a second loss of heterozygosity event rendering it functionally null for BRCA1 protein function. The UWB1.289 BRCA1+ cell line has partial restoration of BRCA1 function from stable expression of a *BRCA1* cDNA construct^61^. We refer to each cell line as BRCA1- and BRCA1+ and together as UWB. IC_50_ values of carboplatin were 36.7μM and 43μM at 72h for the BRCA1- and BRCA1+ cell lines respectively, confirming a previous report **(Supp.** Fig. 7 and Methods) ^62^. Pilot experiments with low (54 μM) and high (180μM) doses of carboplatin showed greater functional responses at the higher dose **(Supp.** Fig. 8). We focused on the latter dose and analyzed 521369 single cells.

### Cell cycle, cell fate and carboplatin uptake responses

In the absence of carboplatin, UWB cells were primarily in G0 and S-phase, with less than 6% in G1 due to TP53 mutations abrogating the G1 checkpoint **(Supp. Fig. 9A, Supp. Table 4)** ^49^. In response to carboplatin, the population of cells in S-phase increased, with BRCA1- cells moving through S-phase more quickly than BRCA1+ cells. By 48hr, 61% of BRCA- cells were in S-phase compared to 88% of BRCA1+ cells. By 72 hr, 42% of BRCA1- cells were still in S-phase, compared to 66% of BRCA1+ cells. This reflects loss of the intra-S-phase checkpoint arrest in BRCA1- cells ^63^. By 72h, 50% BRCA1- cells had entered G0 compared to only 24% of BRCA1+ cells. The number of apoptotic cells increased over time reaching 16% for BRCA+ cells and 15% for BRCA1- cells at 72hr, **(Supp.** Fig. 9B). Comparable intracellular levels of ^195^Pt were detected between genotypes and different fates **(Supp.** Fig. 9B-C).

### Mapping the DDR in single BRCA1- and BRCA1+ cells

Following a similar approach to our single-cell analysis of TYK-nu cells, we clustered the UWB CyTOF data and generated PAGA graphs. In these graphs, clusters of UWB cells with similar DDR profiles formed nodes, and connectivity between these nodes was represented by edges **(Fig. 3, Methods, Fig. 4A–G and Supp.** Fig. 9D and E). UMAP embedding showed a continuum of cells, but with arrangements that were more complex than those observed in TYK-nu cells **(Supp.** Fig. 9D, **Fig. 4A–E)**. To understand the connectivity of subpopulations, we visualized the PAGA graph in a force-directed layout after an additional Leiden clustering on the PAGA graph **(Fig. 5A)**.

**Fig. 5.**
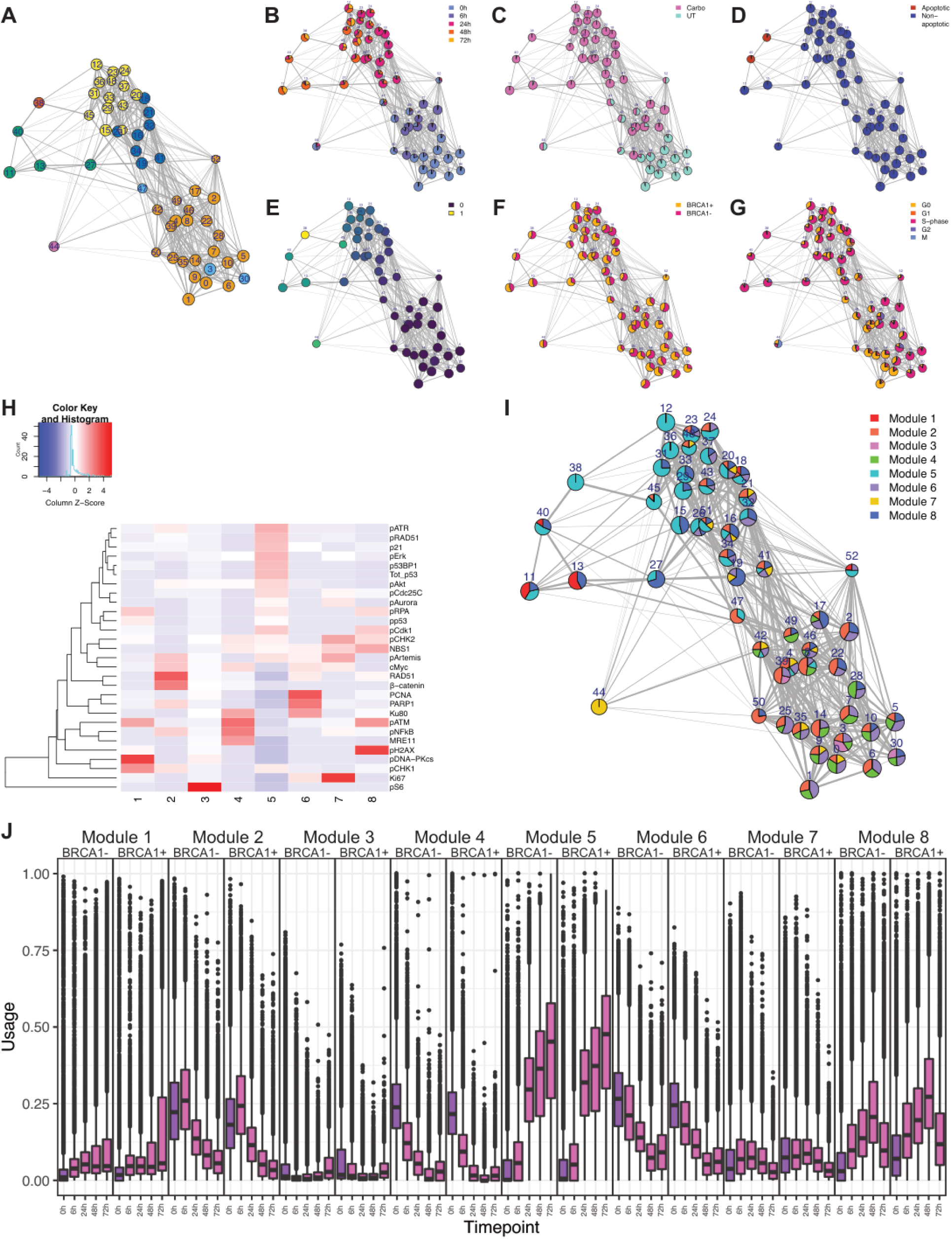
Identification of DDR modules in UWB cells. **A.** PAGA plot of Leiden clusters for UWB cells shows connectivity of clusters in high dimensional space. Cluster nodes are colored based on an additional round of Leiden clustering to identify highly interconnected clusters. **B–G.** PAGA plots colored for time, treatment, cell fate, cell cycle phase, BRCA1-/BRCA1+ and pseudo time. **H.** DDR modules identified by NMF as described in caption **5F**. The contribution of each protein within a module is given by its Z-score depicted on a hierarchically clustered heatmap. **I.** Module activity is depicted by the PAGA plot. Each Leiden cluster is colored with a pie-chart to show the proportion of cells that recruit a specific module. Modules with less than 10% median activity in a cluster were excluded. **J.** Box and whisker plots depict the activity of each module over time.

The PAGA graph for UWB cells revealed a clear separation between both untreated (UT) and cells treated for 6hr compared with their states at 24, 48 after 72hr of treatment **(Fig. 5A–C, Supp.** Fig. 9D). The overlaps between cells at different timepoints observed for TYK-nu cells were mostly absent in the UWB cells. UWB cells treated with carboplatin for 6hrs had a slight shift in their DDR from untreated cells but by 24hr, they had switched to a distinct DDR profile. At 48 and 72hr post treatment, cells followed one of two trajectories with different DDRS but both culminating in apoptosis **(Fig. 5B–D)**. Diffusion pseudo-time (DPT) computed on the PAGA graph confirmed these trajectories **(Fig. 5E)** ^64^. When colored by BRCA1 status, the PAGA graph revealed only minimal separation of cells in late treatment response populations **(Fig. 5F)**. When colored by cell cycle phase, G0 and S-phase cells trended toward different regions of the PAGA graph with M-phase cells separated from the main PAGA graph **(Fig. 5G and I)**. The differences in the timing of DDR responses between TYK-nu and UWB cell lines highlight the need to measure cell states at various stages of treatment.

### Time evolution of DDR in UWB cells revealed by NMF

To determine changes in the DDR of UWB cells over time of carboplatin exposure, we discovered eight DDR modules using NMF. The relative contribution of each protein within a module was depicted with a heatmap **(Fig. 5H)**. Certain proteins that co-occurred in TYK-nu DDR modules also did so in UWB modules, e.g., pH2AX, pATM, and pRPA (Module 8), Myc, RAD51, PCNA, PARP1 (Module 6), Ki67, pChk2, pAurora (Module 7) **(Fig. 5H)**. However, other modules differed between TYK-nu and UWB cells. For example, Module 5, comprised of 13 DDR proteins was not found in TYK-nu cells. Median-module activity in each Leiden cluster was then visualized on the PAGA graph using colored pie charts **(Fig. 5I)**. Both TYK-nu and UWB cells that were in mitosis recruited one module. However, unlike TYK-nu cells, most UWB cells exhibited simultaneous usage of multiple modules. This was especially noticeable in untreated cells, cells 6 and 24hr post carboplatin and cells in G0. All these cells recruited three to five modules but by 48 and 72hr, with a few exceptions, module usage was reduced to one or two.

The temporality of median module usage for individual cells was visualized on box and whisker plots **(Fig. 5J)**. In untreated cells, Modules 2, 4 and 6 were the most active. In response to carboplatin, Module 8, which includes pH2AX, pATM, and pRPA (found in Module 6 in TYK-nu cells) had the greatest activity at 48 hr before decreasing at 72 hours. In contrast, Module 5, the most complex module containing 16 proteins, had significantly greater usage at 24, 48 and 72 hr. This suggests that while Module 8 may be necessary, it may not be sufficient for responsiveness to carboplatin. The complexity of Module 5 suggests that a broader DDR protein network might be needed to promote apoptosis in these highly chemotherapy-resistant cell lines.

### Further characterization of the HGSC DDR landscape in patient-derived cell lines

To ascertain the generalizability of DDR modules, we performed an independent experiment, characterizing the carboplatin-mediated DDR in our recently established CIOV1, CIOV2, CIOV3 cell lines and TYK-nu cells as a control ^41^. These cell lines were spontaneously immortalized continuous lines derived directly from patient tumors. All three cell lines harbored *TP53* mutations and showed varying responses to carboplatin mimicking those in their parent tumors **(Fig. 7A)**. CIOV1 with a non-*BRCA1/2* homologous recombination defective (HRD) phenotype was sensitive to carboplatin. CIOV2 harbored *K-RAS* and *MECOM* amplifications classifying it as innate resistant while CIOV3 harbored a *BRCA1* reversion mutation and was classified as acquired resistant (**Supp. Table 2**). Based on their protein expression levels, the cell lines represented a wide range of HGSC phenotypes reflecting the heterogeneity of primary tumors **(Fig. 6A)** ^38,39^. Specifically, E-cadherin and vimentin delineated cells that were epithelial, mesenchymal, or epithelial-mesenchymal transitioning (EMT) **(Fig. 6A)** ^39^. CIOV1 cells were classified as epithelial because they predominantly expression of E-cadherin. CIOV2 cells displayed a mix of epithelial and EMT phenotypes, with some cells expressing E-cadherin alone whereas others co-expressed E-cadherin and vimentin. In contrast, CIOV3 cells were primarily mesenchymal, with a small subset showing characteristics of EMT.

**Fig. 6.**
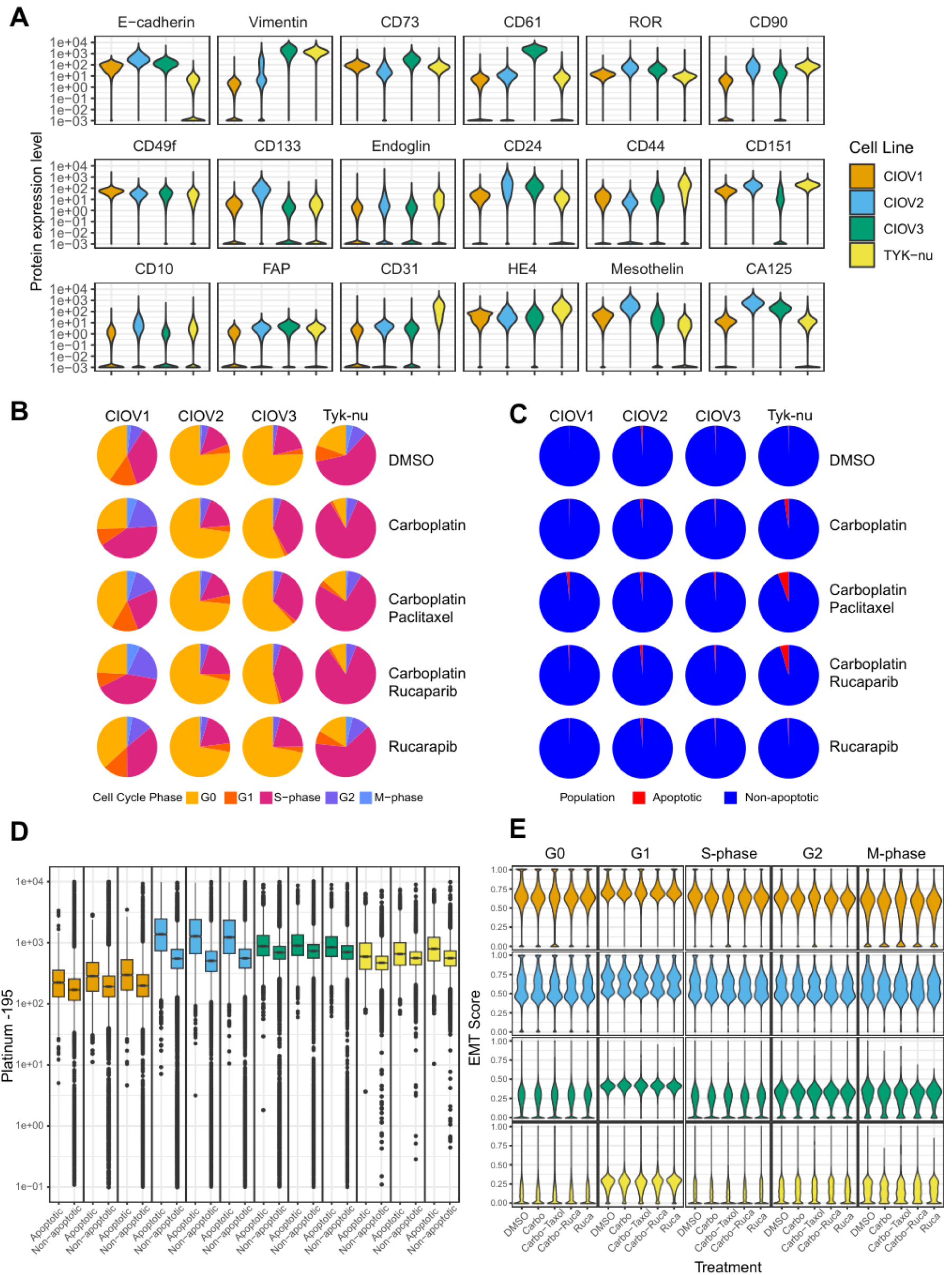
Characteristion of patient-derived CIOV1–3 cell lines. **A.** Violin plots depicting expression of epithelial, mesenchymal, stem cell, and HGSC proteins. Key colored for cell line. **B.** Cell cycle distributions vary across cell lines in response to treatments. **C.** Pie charts showing minimal apoptosis under the conditions chosen. **D.** Box and whisker plots depicting 195-Pt uptake. Box colors correspond to cell lines as in in Fig. 6A. Left to right within each cell line are treatment with carboplatin, carboplatin + paclitaxel, and carbplatin +rucaparib. **E.** Violin plots depicting changes in epithelial and mesenchymal states within each cell cycle phase in response to treatment. EMT scores range from 0 to 1 with score of 1 indicating a purely epithelial phenotype and a score of 0 indicating a purely mesenchymal phenotype and defined by levels of E-cadherin and Vimentin.

**Fig. 7.**
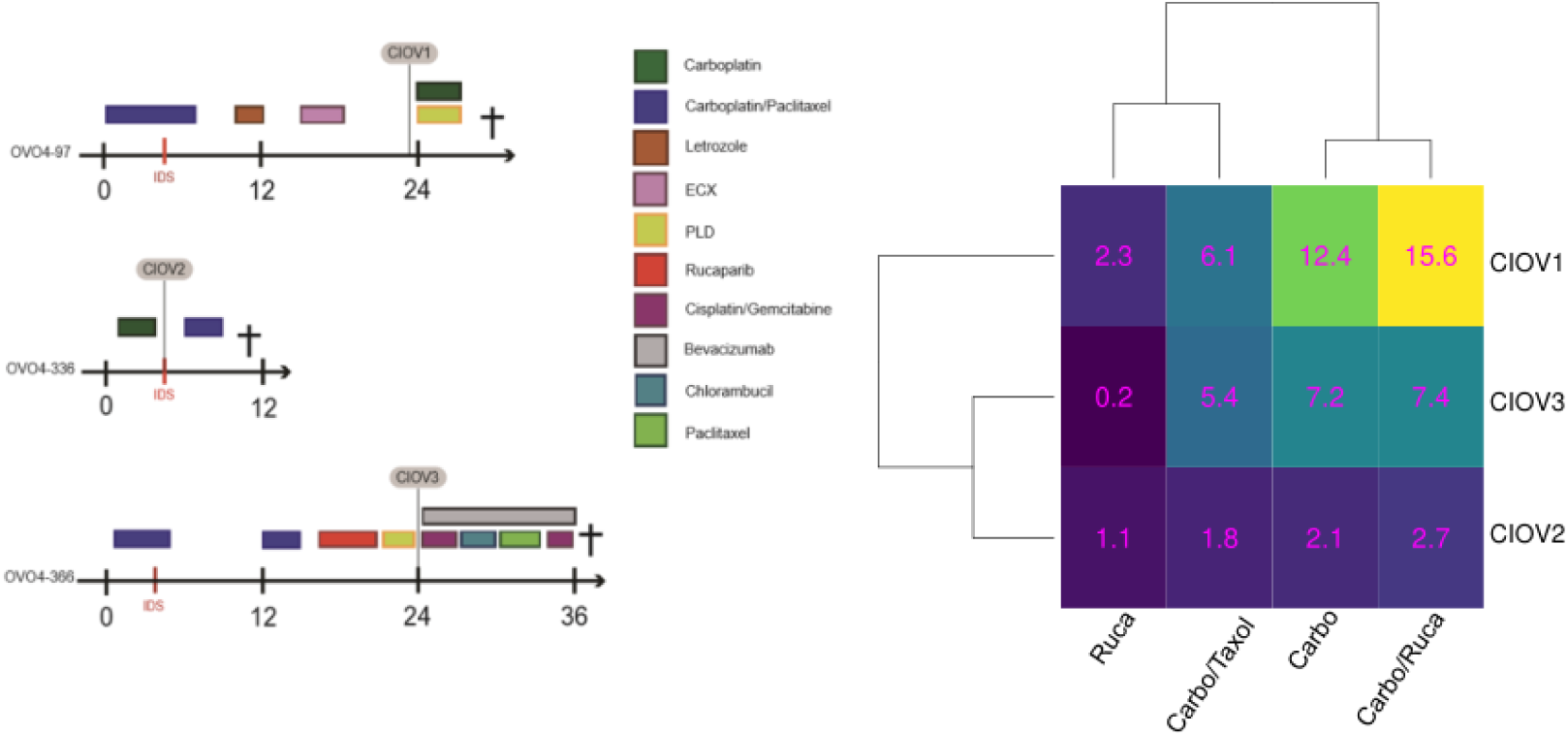
Validation of TYK-nu NMF Module 6 in CIOV cell lines. In an independent experiment, CIOV1, 2 and 3 cell lines were treated with a carboplatin-based regiment for 48 hr, and processed for CyTOF with the same DDR antibody panel. **A.** Treatment journey showing patients whose tumors were responsive or resistant to chemotherapy. Time for tumor acquisition is shown when samples were placed into 2D cell culture to derive CIOVs 1, 2 and 3. **B.** Hierarchically clustered heatmap depicting TYK-nu NMF Module usage. The values in the heatmap are pecentage increase in module activity relative to DMSO control in CIOV cell lines under conditions shown.

Cell lines were treated with carboplatin (8 μM), carboplatin (8 μM) + paclitaxel (5 nM), carboplatin (8 μM) + rucaparib (1.2 μM), rucaparib (1.2 μM), or DMSO (control) for 48hr. To mimic a clinical setting the CIOV cell lines all received the same drug doses. At 48 hr cells were harvested, bar-coded, combined, stained with the antibody panel and processed for CyTOF producing a dataset of 7,206,826 cells for downstream analysis **(Fig. 1B, Supp. Table 1** and **Methods)**.

### Cell cycle phase, cell fate, and carboplatin uptake in CIOV cell lines

The cell lines showed different cell cycle phase distributions at baseline and in response to treatments **(Fig. 6B)**. Carboplatin alone led to varying levels of cells in S-phase. Adding paclitaxel or rucaparib to carboplatin produced varying effects on other cell cycle phases. Paclitaxel increased cells in G0 for CIOV1, but in G0 and G2 for CIOV2 and TYK-nu. Rucaparib combined with carboplatin induced minimal effects in all cell lines. With its innate resistant phenotype, CIOV2 was least affected by all treatments. Under the drug treatment conditions studied, most cells survived with minimal apoptosis **(Fig. 6C)**.

There was considerable variability in carboplatin uptake among individual cells which was not affected by adding paclitaxel or rucaparib. While the interquartile ranges (IQRs) showed significant overlap between apoptotic and non-apoptotic cells in CIOV1 and CIOV3, there was much less overlap in CIOV2. This suggests that drug efflux potentially through upregulated transporters may have a greater role in the innate resistant CIOV2 cell line **(Fig. 6D)**.

### Relationship between epithelial mesenchymal phenotypes with cell cycle phase

Given the emergence of drug resistance in epithelial/mesenchymal transitioning cells, the relationships between epithelial/mesenchymal states, cell cycle and treatments across the CIOV cell lines were determined ^65^. As a proxy for epithelial/mesenchymal phenotype, we computed an EMT score using levels of E-cadherin and vimentin in single cells. An EMT score of 1 implies an epithelial phenotype while a score of 0 implies a mesenchymal phenotype **(Fig. 6E and Methods)**. CIOV1 cells were the most epithelial in all cell cycle phases, while CIOV2 cells were comprised of a mix of phenotypes. CIOV3 cells more closely mirrored TYK-nu cells which were previously shown to be mesenchymal ^39,66^. EMT scores changed marginally in response to treatments but significantly within a cell cycle phase. The cell lines trended toward an epithelial phenotype in G1, and toward a mesenchymal phenotype in the other cell cycle phases, a result, to the best of our knowledge, not reported previously. The drug resistance in CIOV2 (innate) and CIOV3 (acquired) is consistent with the presence of cells with EMT and mesenchymal phenotypes ^67^.

### NMF module activity associated with clinical response

Given that the CIOV cell lines closely resembled their parent tumors in genetic, molecular, and chemotherapy responses, we investigated if any NMF modules tracked with *in vivo* and *in vitro* chemotherapy responses **(Fig. 7)**. We analyzed the CIOV1-3 CyTOF datasets using NMF Module 6, which had the highest activity in TYK-nu cells at late exposure times to carboplatin **(Fig. 4H)**. Compared to vehicle-treated cells, Module 6 was most active in CIOV1 cells, minimally active in CIOV2, consistent with its innate resistance, while there was intermediate activity in CIOV3. Module 6 usage was greater for the combination of carboplatin with rucaparib compared with standard-of-care paclitaxel for all three CIOV cell lines. By contrast, Module 5, discovered in UWB cells, had minimal usage in all CIOV cell lines. These findings support the use of Module 6 recruitment as a potential indicator of response to chemotherapy.

## Discussion

Our study of carboplatin resistance in human ovarian cancer was predicated on the hypothesis that in response to carboplatin tumor cells selectively activate specific DDR protein sub-networks or modules through changes in phosphorylation and abundance. Activating the entire DDR network would be metabolically inefficient. To understand the functional consequences of genetic, transcriptomic and epigenetic changes which result in sensitivity or resistance to carboplatin, we capitalized on the single cell attributes of CyTOF. to measure the carboplatin-mediated DDR. To understand the different fates of cells with comparable levels of intranuclear carboplatin, we analyzed the CyTOF datasets with UMAP embedding, PAGA and NMF **(Fig. 3)**. UMAPs generated from intracellular DDR, cell cycle and signaling proteins revealed a continuum of cells exhibiting a range from subtle to large differences in their DDR profiles **(Fig. 4)**. These observations align with a recent study showing that ovarian tumor cells transition through a series of transcriptomic states as they progress toward resistance to a PARPi ^25^.

Applying NMF we showed that DDR(s) in individual cells can be characterized by distinct protein modules. TYK-nu cells tended to use one DDR protein module at a time, regardless of conditions, such as cell cycle phase or duration of treatment. By contrast, highly resistant UWB cell populations used multiple modules simultaneously. This was most evident for cells that were untreated or exposed to carboplatin for 6 or 24 hr **(Fig. 5)**. It could be that after the patient received multiple rounds of chemotherapy, the cells had reached a state of full resistance potentially maintained by recruitment of multiple DDR protein modules. Nevertheless, in M-phase, both TYK-nu and UWB cells recruited one unique module that contained Ki67 and pAurora, two proteins with established roles in M-phase ^68,69^. Stable reintroduction of *BRCA1* into UWB-*BRCA1*-null cells made little difference to their overall module usage or resistance to carboplatin. While reversion mutants of *BRCA1* and *BRCA2* confer resistance by restoring HR, in this case introduction of *BRCA1* had minimal effect on a tumor cell that was likely maximally resistant ^17,20^.

For TYK-nu and UWB cells, modules containing the transcription factors pNFKb, pMyc and β-catenin (Module 3 in TYK-nu and Module 2 in UWB) were active at early times. These transcription factors have established roles in promoting chemoresistance, stemness and survival ^70–72^. Modules containing pRPA, pATM, pH2AX and pCHK1 or pCHK2 (Module 6 in TYK-nu and Module 8 in UWB) indicate active DNA repair and cell cycle arrest ^73^. In TYK-nu cells, Module 6 usage occurred at late times when cells were in pre- and early apoptotic states suggesting that DNA repair efforts had failed **(Fig. 4G, H)**. In UWB cells, recruitment of Module 8 was replaced by increased reliance at later time points (24, 48 and 72 hours) on the more complex Module 5 **(Fig. 5I, J)**. This suggested that a larger DDR protein module was necessary to maintain therapeutic resistance consistent with the transcriptomic complexity as described ^24,25^.

To validate our findings, we interrogated our recently generated CIOV cell lines for their usage of Module 6 discovered in the pre-apoptotic TYK-nu cells. The presence of EMT and mesenchymal cell phenotypes in CIOV2 and CIOV3 is consistent with their resistant characteristics ^65^. Unexpectedly, we observed cell cycle -dependent changes in the EMT score with more epithelial phenotypes for cells in G1. This suggests a mechanism whereby E-cadherin, through its cell adhesive function with β-catenin may regulate levels of transcriptionally active β-catenin and consequently downstream target genes associated with proliferation such as *MYC* and *CYCLIN D1* ^74^. In contrast, the shift toward a more mesenchymal phenotype during S-phase and G2 may enable cells to over- ride cell cycle checkpoint arrest and adapt to carboplatin-mediated DNA damage ^65^. Module 6 usage was linked to clinical responsiveness to carboplatin across the three CIOV cell lines. It was highest in carboplatin-sensitive tumor cells (CIOV1), lowest in those with innate resistance (CIOV2) and intermediate in cells with acquired resistance (CIOV3). Cells treated with carboplatin plus rucaparib showed the highest usage of Module 6, while carboplatin plus paclitaxel showed the lowest. This was most marked for CIOV1 perhaps because this patient had not received rucaparib during their treatment. The minimal usage of the complex UWB Module 5 **(Fig. 5H)** across the CIOV cell lines (data not shown) suggests recruitment of alternate protein modules. This may reflect both greater complexity and plasticity required to maintain carboplatin resistance ^75^.

This study is limited by the inherent complexity of the DDR. Our CyTOF panel did not provide full coverage of the DDR, thus potentially missing additional DDR modules. Nevertheless, it successfully measured 36 phosphorylation states and protein levels with established roles in the DDR, cell cycle and signaling ^73,76,77^. While interactions with immune cells and stroma within the tumor microenvironment were not explored, focusing on tumor cell autonomous mechanisms is a critical first step in unravelling DDR complexity linked to carboplatin resistance.

Many of a tumor’s adaptive responses to therapy are targetable ^29^. However, differences in innate sensitivity to carboplatin between individuals and acquired resistance make it difficult to determine which adaptive pathway(s) to target and the optimal timing during a patient’s treatment journey. We propose that monitoring Module 6 usage as a readout in drug screens of carboplatin combined with other medicines in CIOVs could identify more beneficial therapeutic combinations for patients. The different resistance states of CIOVs make them a valuable resource toward this endeavor.

Furthermore, our cell suspension CyTOF assays can be readily adapted for spatial analyses allowing for broader characterization of resistance modules particularly in in vitro model systems. This will refine the selection of antibody panels reading out both DDR and immune responses in clinical trial samples. Our approach is generalizable to study drug resistance in other types of cancer.

## Methods

### Cell lines

HeLa and OVCA3 (American Type Culture Collection (ATCC)) and TYK-nu (National Institute of Biomedical Innovation, Japanese Collection of Research Bioresources Cell Bank (JCRB)) cell lines were cultured in Dulbecco’s Modified Eagle Medium (DMEM) (Gibco), McCoy’s 5A modified medium (Gibco) and Eagle’s minimal essential medium (EMEM) (ATCC) respectively supplemented with 10% heat-inactivated fetal bovine serum (FBS) (Hyclone), 1% Penicillin-Streptomycin (Gibco) and 2 mM L-glutamine (Gibco). UWB1.289 -/-BRCA1 and +BRCA1 cell lines (ATCC) (referred to in the main manuscript a BRCA1- and BRCA1+) were cultured in 50% RPMI-1640 (Gibco) supplemented with 2mM L-glutamine, 25 mM sodium bicarbonate and 50% mammary epithelial growth medium (MEGM) (Lonza) supplemented with 3% heat-inactivated fetal bovine serum (FBS). G-418 (200 mg/mL Geneticin from Gibco) was added to the media for the UWB1.289 BRCA+ cell line. JHOS2 (RIKEN BRC Cell Bank) cell line was cultured in Dulbecco’s Modified Eagle Medium/Nutrient Mixture F-12 (DMEM/F-12 (Ham), Gibco) supplemented with 10% FBS, 0.1 mM MEM non-essential amino acids (NEAA, Gibco) and 1% penicillin/streptomycin. CIOV1, 2, 3 cell lines (Brenton Lab, University of Cambridge, UK) were cultured in DMEM/F-12 medium (Gibco) with 10% FBS and 1% penicillin/streptomycin. Cells were split every 2–3 days and kept in a humidified cell culture incubator at 37°C with an atmosphere of 95% air and 5% CO_2_.

### Dose response curves for carboplatin, talazoparib, rucaparib and paclitaxel

Carboplatin (Sigma Aldrich) was dissolved in water (13.5 mM stock solution) and talazoparib (MedChem Express) was dissolved in DMSO (10 mM stock solution). Cells were seeded in 96-well flat-bottomed plates (5000 TYK-nu cells per well for carboplatin treatment, 2000 TYK-nu cells per well for talazoparib treatment, 2000 TYK-nu cells per well for carboplatin plus talazoparib and 2000 BRCA1- and BRCA1+ cells per well for carboplatin treatment). For measuring the IC_50_ values for each drug alone, serial dilutions were performed. Starting concentrations were 1.6 mM for carboplatin and 10μM for talazoparib. TYK-nu cells were exposed continuously to each drug in a two- or three-fold serial dilution (9 or 11 points) for 24, 48, 72h or 96 h in triplicate and cell growth inhibitory effects were determined using the Vybrant® MTT Cell Proliferation Assay (ThermoFisher Scientific) **(Supp.** Fig. 3A, B). We subsequently transitioned to the RealTime-Glo™ MT Cell Viability Assay (Promega) which has the convenience of monitoring cell viability continuously using the same sample well generating more information about drug action with respect to time and dose dependence. We used this assay, according to the manufacturer’s recommendations, to determine the IC_50_ concentrations for carboplatin and talazoparib for cells treated with the combination. Cells were exposed to three-fold fixed ratio serial dilutions of a starting mixture of carboplatin (16μM) and talazoparib (3 μM). Luminescence was measured every 24 h until 72 h **(Supp.** Fig. 3C and 3D). We tried several pilot experiments holding the concentration of each drug constant and varying the other (data not shown). However, the conditions chosen, 8μM carboplatin and 100nM for talazoparib were optimal for keeping enough cells viable for CyTOF assays particularly at the later timepoints. Lower drug concentrations had only subtle effects on cell cycle/DDR measurements for the times chosen. The RealTime-Glo™ MT Cell Viability Assay was also used to determine carboplatin IC_50_ values for BRCA1- and BRCA1+ cells **(Supp.** Fig. 7). The IC_50_ values at 72 h were used to guide the carboplatin concentrations chosen for experiments, 54 and 189**μ**M.

### Cell line treatments

HeLa cells were cultured in 10cm dishes to a confluency of ∼80%. They were exposed to the following DNA damaging agents: 5Gy ionizing radiation (IR) **using a Cesium 137 irradiator,** followed by a 30 min rest, 100 J/m^2^ ultraviolet C (UVC) followed by a 90min rest, 10μM etoposide (Sigma Aldrich) with 0.02% DMSO control for 24h or 316.7μM carboplatin for 24 or 48h with H_2_O as control. Cells were also treated with 100ng/mL nocodazole (Sigma Aldrich), a microtubule inhibitor for 18h with 0.01% DMSO control.

TYK-nu cells were treated with 8μM carboplatin or 8μM carboplatin plus 100nM talazoparib for 24, 48, 72 and 96h, with H_2_O as control harvested at 96 h.

UWB.289 BRCA1- and BRCA1+ isogenic cell lines were treated with 54μM or 180μM carboplatin for 6, 24, 48 and 72h with H_2_O as a control for 72 hr. CIOV1, CIOV2, CIOV3, and TYK-nu cells were treated with 8μM carboplatin, 8μM carboplatin + 5nM paclitaxel (Sigma-Aldrich, dissolved in DMSO), 8μM carboplatin + 1.2μM rucaparib (Selleckchem, dissolved in DMSO), or 1.2μM rucaparib for 48h with 0.012% DMSO control for each cell lines. Treatment conditions for cell line experiment were performed in triplicate. Each time after treatment, cells were incubated with 10mM iododeoxyuridine (IdU) (Sigma Aldrich) and viability dyes. For HeLa cells treated with all agents, aside from carboplatin, cells were treated with 25mM cisplatin (Sigma Aldrich) for 1 min as previously described ^34^. For experiments with carboplatin, cells were treated with 1mM Cell-ID™ Intercalator- 103Rh (Standard BioTools) for 15 min as previously described ^34^. Cells were subsequently harvested, fixed with 1.6% paraformaldehyde, washed twice with CSM, flash-frozen, and stored at - 80°C ^34,39^.

### Isolation of nuclei

Untreated and carboplatin-treated 4–5 x 10^6^ TYK-nu cells were harvested at different timepoints (24, 48, 72, and 96 h). Half the cell aliquot was processed as described above. The other half of the aliquot was processed for isolating nuclei. TYK-nu cells, cell pellets were resuspended in 300 µL of cold lysis buffer (Tris-HCl pH 7.4 (10 mM), NaCl (10 mM), MgCl_2_ (3 mM) and Igepal CA-630 (0.025% in PBS, Millipore Sigma) and incubated on ice for 10 min (time optimized from 10X Genomics protocol). The reaction was quenched with 1.2 mL of cold cell staining media (CSM) and suspensions of nuclei were washed twice with CSM (500 x g, 10 min, 4°C). Nuclei were fixed in 1 mL 1.6% PFA in PBS for 10 min at room temperature, washed twice with CSM, resuspended in CSM (∼150-200 µL), snap frozen in dry ice, and stored at 80°C.

### Confirming purity of isolated nuclei

The purity of the nuclei was evaluated using an anti-histone H3 antibody (D1H2 (Standard BioTools)) by CyTOF and then gating for non-apoptotic and apoptotic cells using a c- PARP antibody **(Supp.** Fig. 4A). Additionally, purity of the nuclei was determined by comparing their scatter properties with those of whole cells **(Supp.** Fig. 4B) using an LSR2 flow cytometer. Further confirmation of purity was confirmed by microscopy with DAPI (blue) and vimentin-Ax647 (D21H3 (CST)) using a Keyence BZ-X800 microscope **(Supp.** Fig. 4C).

### Antibodies for CyTOF

Antibodies were all conjugated in-house **(Supp. Table 1)**. In brief, antibodies in carrier-free PBS were conjugated to metal-chelated polymers (MaxPAR antibody conjugation kit, Standard BioTools) according to the manufacturer’s protocol. Bismuth-chelated polymer labeling was performed with an in house- protocol ^78^. Metal-labeled antibodies were diluted to 0.2–0.4 mg/mL in antibody stabilization solution (CANDOR Biosciences) and stored at 4°C. Each antibody was titrated using positive and negative controls as described **(Supp. Table 1)**. Antibody concentrations chosen were based on optimal signal-to-noise ratio.

### Sample processing and antibody staining for CyTOF

Frozen, fixed single-cell suspensions of cell lines were thawed at room temperature. For each sample, 1 x 10^6^ cells were aliquoted into cluster tubes in 96 well plates and subjected to pre-permeabilization palladium barcoding ^79,80^. After barcoding, cells were pooled, washed, and incubated for 5 min at room temperature with FcX block (Biolegend,) to block non-specific antibody binding. Cells were then incubated with the CyTOF antibody panel, washed, and incubated with the ^191/193^Ir-intercalator at 4°C overnight. Cells were resuspended in a solution of normalization beads washed and resuspended before introduction into the CyTOF2 ^39^.

### Sample processing and antibody staining of isolated nuclei

Fixed frozen TYK-nu nuclei were thawed at room temperature. Samples were transferred into cluster tubes containing 1 mL of cold CSM and washed (600 x g, 10 min, 4 °C). Samples were permeabilized in 1 mL 100% ice-cold methanol for 20 min at 4 °C, washed twice with cold CSM and stained with antibodies against vimentin (D21H3 (CST)) and intra-nuclear markers (Histone H3, c-PARP (F21-852 (BD)) for 1 h at room temperature on a shaker. Samples were washed twice with cold CSM and incubated in 1 mL ^191/193^Ir DNA intercalator solution (0.1 µM) in 1.6% PFA (PBS) overnight at 4 °C. TYK-nu nuclei suspensions were washed once with CSM and twice with CyTOF water, prior being resuspended in a solution of normalization beads and introduced into the CyTOF2. Platinum was read out on the 195 channel, as it represents the most abundant stable platinum isotope.

### Processing frozen nuclei for microscopy

Fixed frozen TYK-nu nuclei were thawed at room temperature, transferred to FACS tubes containing 1 mL of cold CSM and washed (500 x g, 10 min, 4 °C). Samples were permeabilized in 100% ice-cold methanol (1 mL) for 20 min at 4 °C, washed twice with cold CSM and stained with vimentin-A647antibody (5 µL in 100 µL of reaction) for 30 min on ice. After addition of 2 mL CSM, samples were washed twice with CSM followed by staining with DAPI (1 µg/ mL in CSM, 500 µL) for 10 min at room temperature. After two washes with CSM, nuclei were resuspended in 100 µL of CSM. 10 µL of nuclei in suspension were transferred onto a microscope slide, a coverslip was placed on top samples were imaged using the Keyence microscope BZ-X800.

### Data analysis tools and illustration design software

All data, statistical analysis, and figures were conducted with Adobe Illustrator, Microsoft Excel, Microsoft PowerPoint, R 4.1.2, Python 3.7, MATLAB 2019, and GraphPad Prism 8. CyTOF datasets were evaluated with software available from Cytobank and CellEngine. The study schematic and signaling map (Fig. 1) were created with BioRender.com. Biaxial plots **(Fig. 2B)** were generated in CellEngine (https://cellengine.com). Dose response curves and IC-50 values in **Supp. Fig. 3C, Supp., Fig 4**, and **Supp. Fig. 7** were generated using GraphPad Prism 8. Supplementary Figure 4A, 4B, and 5A were generated with Microsoft PowerPoint. Multiplexed Louvain community detection in Fig 4C was performed using a modified script in MATLAB 2019 http://netwiki.amath.unc.edu/GenLouvain, https://github.com/GenLouvain/GenLouvain (2011-2019). and a custom pre-processing script written in R. All other analyses and figures were generated using custom R and Python scripts written in-house that are publicly available. Specific package requirements for scripts are included in code. Analyses in CellEngine and Cytobank were performed at cellengine.com and cytobank.org, respectively. Analyses in Python and MATLAB were performed on a custom-built server running Windows 10 with 256 GB RAM. All other analyses were performed on a MacBook Pro with 64 GB RAM.

### Initial assessment of data quality and cell fate identification

CyTOF FCS files were normalized and debarcoded using algorithms reported previously ^80,81^ with access at the two links https://github.com/ParkerICI/premessa

https://github.com/nolanlab/bead-normalization/wiki/Normalizing-FCS-Files

Tailored manual gating was performed in the Cytobank or CellEngine software. Singlets were gated based on ^191^Ir/^193^Ir DNA content and event length to exclude debris and doublets. Following singlets gating, cells were gated using viability dye (^103^Rh or cisplatin) into dye positive and dye negative populations. Dye negative populations were further gated based on levels of c-PARP into non-apoptotic/viable (c-PARP-) and apoptotic (c- PARP+). Cisplatin was used as a viability dye in HeLa cells ^35^. For experiments with carboplatin treatments, ^103^Rh was used as an alternate viability dye.

### Measurements of cell cycle

Cell cycle distribution was measured by applying the manual gating strategy described previously ^37^. Viability dye negative cells were used to analyze the cell cycle for each condition. Gating strategy is summarized in **Supp. Fig. 1**. To delineate cell cycle phases, we first utilized IdU to identify cells in S phase, and an antibody against cyclin B1 to demarcate the rest of the cells in G0/G1 and G2/M phases. Antibodies against pRb (S807/811) and cyclin E were applied to separate G0 and G1 phases. G2 and M phases of the cell cycle were distinguished by gating with antibody against pHH3 (S28).

### Cell Line DDR Mutation Profiles

GO analysis was performed using AmiGO2 searching Genes and Gene Products with keyword DNA+Damage+Response. From organism drop down Homo sapiens was selected and from Type column protein was selected. Results were downloaded as txt file on April 28, 2022. Results from depmap were downloaded by searching for cell lines from Cell Line Selector. Mutations were downloaded. Then script in R was used to match genes identified with GO analysis to mutated genes in cell lines and results were saved in **Supp. Table 2**.

### Computational analysis Cell cycle phase pie charts

Cell cycle phase pie charts were computed as the proportion of non-apoptotic cells in each cell cycle phase and generated in ggplot2.

### Protein expression violin plots

Violin plots were generated in ggplot2 using live cells (Cisplatin-negative for HeLa cells in Figure 2 and Rh-103 negative for all other single cell data)

### Platinum uptake box and whisker plots

Notched box and whisker plots were generated in ggplot2. The notches extend 1.58 * IQR / sqrt(n) which gives a roughly 95% confidence interval for comparing medians [REF McGill et al. (1978)]. Data were log_10_ normalized prior to visualization.

### Fold change nuclei isolation

Median platinum levels were extracted for both non-apoptotic and apoptotic cells and nuclei. The ratio of median nuclear to cellular platinum levels for both apoptotic and non-apoptotic cells was used to construct the bar chart.

### LDA

Linear discriminant analysis (LDA) was computed using the MASS package in R. A training set and test set with balanced classes were generated from viable HeLa single cell data. For each treatment (class), 7000 cells were randomly sampled then randomly partitioned into 6300 cells for the training set and 700 cells for the test set. This resulted in a training set of 31500 cells and a test set of 3500 cells. Linear discriminant functions (LDFs) were fit using the MASS::lda function on the single cell DDR protein expression levels for cells in the training set. Markers for cell cycle phase, apoptosis, and viability were excluded. To test the quality of fit, four LDFs were used to predict treatment from DDR protein levels on the test set and classification results were reported in confusion matrix. The macro-F1 score was used as a summary statistic for the performance of the final fit LDFs. The macro-F1 score is defined as the average F1 score over all treatments and is summarized in the following equation:

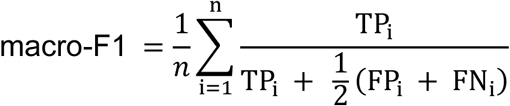

where the value in the sum is the F1 score for treatment *i* and TP_i_, FP_i_, and FN_i_ denote the number of true positives, false positives, and false negatives for group *i*, respectively. The waterfall plot displaying the loadings of LDF1 was generated using ggplot2 in R.

### Single cell data visualization

Single cell data were visualized using the scanpy package in Python ^82^. To denoise the data, cells with total expression (sum over all marker expression levels) higher than the 99.5% quantile were excluded. For normalization, we apply the approach to the standard normalization pipeline in Monocle3 (https://cole-trapnell-lab.github.io/monocle3). After normalization, principal component analysis (PCA) was applied to reduce the data dimensionality. Based on a waterfall plot of % variance explained vs. principal component (PC), the top 10 PCs with highest % variance explained were selected. Next, a nearest neighbor graph was constructed on the single cell data, with k = 30 nearest neighbors. Uniform Manifold Approximation and Projection (UMAP) was applied to this data with default parameters and the results were visualized using ggplot2. This procedure was performed separately for HeLa cells **(Supp.** Fig. 2D, E**, Supp.** Fig 3D-G), TYK-nu cells **(Fig. 4A)**, and UWB cells **(Supp.** Fig. 9D).

### Clustering, PAGA, and DPT

Single cell data were clustered with the Leiden algorithm in scanpy ^57^. The Leiden algorithm was selected since it has been shown to identify connected communities more accurately than Louvain. The role of clustering in this study is not to identify distinct cell phenotypes but to define similar (often overlapping) groups of cells whose characteristics (such as cell cycle phase, treatment, and cell fate) can be analyzed in the context of the larger dataset. In this respect, the goal was to generate a sufficient number of clusters to cover the manifold while keeping the number low enough to enable reasonable visualization and downstream analyses. Clustering was performed separately for the TYK-nu and UWB time course experiments. For both datasets, pre-processed as described in the previous section, a nearest neighbor with k = 30 nearest neighbors was constructed, and Leiden clustering with resolution = 5.0 was run. Results were visualized in ggplot2. After clustering, PAGA was used to compute the connectivity between the identified clusters with default parameters ^58^. Edges with PAGA weights less than 0.1 were excluded. PAGA graphs were visualized in the R igraph package. Diffusion pseudotime (DPT) **(Fig. 5E)** was computed with default parameters using the scanpy package.

### NMF

NMF was computed using the consensus NMF approach with the scikit-learn package in Python ^59^. To denoise the data, cells with total expression (sum over all marker expression levels) higher than the 99.5% quantile were excluded. Data were then row normalized and scaled to unit variance. Consensus NMF modules were then computed by computing 100 sets of modules using the NMF function in scikit-learn, with max_iter = 3000. K was set to 30 for KNN-deviation and only modules that fall within a distance threshold of 0.1 were kept. Resulting modules were visualized using the gplots package in R and ComplexHeatmap **(Figs 4F and 5H)**. For cell line specific NMF, eight modules were computed using the above approach for each cell line separately. Box and whisker plots showing module activity over time were generated in ggplot2.

### Resource availability Lead contact

Further information and requests for resources and reagents should be directed to and will be fulfilled by the lead contact, Wendy J. Fantl

## Materials availability

This study did not generate new unique reagents.

## Data and code availability

All data reported in this paper will be shared by the lead contact upon request.

## Acknowledgements

This work was supported by funding from the BRCA Foundation and the V Foundation for Cancer Research; a gift from the Gray Foundation, Department of Defense (W81XWH-12-1-0591), NCI (1R01CA234553, R21CA231280), the 2019 Cancer Innovation Award, the 2021 Cancer Innovation award both supported by the Stanford Cancer Institute, an NCI-designated Comprehensive Cancer Center, the Department of Urology, Stanford University; NHLBI (P01HL10879709); NIAID (U19AI057229); and a PICI Bedside to Bench grant. A.D.-G. thanks the Fundacion Alfonso Martin Escudero and Ovarian Cancer Research Alliance for Mentored Investigator Award (MIG-2023-2-1015) for his postdoctoral fellowships. We wish to thank Dr Zach Bjornson for his design of new software in CellEngine to enable part of our data analysis. We wish to thank Dr. Keith Shults and others for critical reading of the manuscript and Professor Garry Nolan for the use of the CyTOF2 mass cytometer.

## Author contributions

Conceptualization, W.J.F; J.D.B, J.S.S Methodology, V.D.G, Y.W-H, M.V, A.D.G, Validation, J.S.B, I-G.F, Investigation, V.D.G, Y.W-H, A.D.G, Resources, V.D.G, Y.W-H, M.V, A.D.G; Formal analysis, J.S.B, Z.R, A.M, A.L Data curation, J.S.B, Writing – Original Draft, J.S.B; W.J.F Writing – Review & Editing, W.J.F, J.D.B, A.A, J.S.B I-G.F Visualization, J.S.B, I-G.F, A.D.G, W.J.F, Funding Acquisition, W.J.F, A.A, J.D.B Resources, M.E.V and C.K.B.; Supervision, W.J.F, J.D.B, A.A., Project administration, W.J.F.

