## Supplementary Text, Figures and Tables for "Coordinated protein modules define DNA damage responses to carboplatin at single cell resolution in human ovarian carcinoma models"

##### Validation of CyTOF panel using Hela cells exposed to genotoxic agents and a microtubule inhibitor

To validate our CyTOF antibody panel for studying the DDR in single cells, we first performed experiments in asynchronously growing HeLa cells, as this model has been extensively characterized for DDR responses to multiple agents (ionizing radiation (IR), ultraviolet radiation C (UVC), etoposide or nocodazole [1-3] (**Supp. Fig. 2)**. After treatment, cells were incubated with iododeoxyuridine IdU, and cisplatin, barcoded with palladium isotopes, pooled, stained with the antibody panel, and processed for CyTOF [4-8]. Single cell datasets were subject to quality control before analysis **(Methods)**.

As expected, the cell cycle response differed for each treatment. Compared to untreated cells, low dose IR induced a modest accumulation of cells in G1 and S, while UVC induced a modest G1 arrest **(Supp. Fig. 2A)**. Etoposide, a topoisomerase II inhibitor, and inducer of double-strand breaks (DSBs) incurred a three-fold accumulation of cells in S-phase and G2, whereas cells treated with nocodazole, a microtubule inhibitor, accumulated in G2/M [2]. While these cell cycle measurements confirmed established results, they did not provide insight about the relationship between cell cycle phase and the DDR which necessarily differs between treatments [9-12].

##### Genotoxic agents mediate distinct changes in DDR protein expression levels in HeLa cells

DDR pathways are known to be selectively engaged in different phases of the cell cycle However, for each treatment, CyTOF revealed significant variability of DDR protein expression levels across each cell cycle phase, suggesting that individual cells can undergo a wide spectrum of responses **(Supp. Figs. 2B)**. To explore how individual proteins participate in these diverse DDRs, we first identified those proteins with the greatest differences in expression levels between treatments by performing linear discriminant analysis (LDA), a dimensionality reduction algorithm and classifier, on the single cell HeLa protein expression data from the five conditions **(Methods, Supp. Fig. 2C)** [20]. LDA computed a ranked set of linear discriminant functions (LDFs) that optimally separated the datasets, with LDF1 being the most discriminating. The LDFs accurately predicted treatment from protein expression levels on a test set of single cell HeLa data, as shown by a macro-F1 score of 0.84 **(Methods and Supp. Fig. 3A)**. For LDF1, pNFkB had the highest positive weight and pH2AX and pATM the highest negative weights **(Supp. Fig. 2C)**. This is consistent with their known biological activation in response to DSBs, and the potency of different perturbations to induce DNA DSBs; low for UT and nocodazole, and moderate and high for UVC, IR and etoposide [13-15].

Median expression levels of pNFkB, pH2AX and pATM were also consistent with the directionality of the LDF1 weights, confirming the utility of the LDA **(Supp. Fig. 3B)**. These findings confirmed that our antibody panel detected specific DDRs for specific treatments and secondly, that large scale data analysis methods can identify the proteins driving different treatment responses.

**Deconvoluting relationships in Hela cells between DDR and cell cycle in response to treatments**

To gain further insight into the complex interplay in individual cells between treatment, cell cycle phase, and the DDR, we visualized our single cell data using Uniform

Manifold Approximation and Projection (UMAP) **(Supp. Fig. 2D, 2E)** [16]**.** We excluded proteins for cell cycle phase and apoptosis to generate UMAPs based exclusively on DDR and cell cycle regulatory proteins. This analysis revealed three key observations. First, some DDRs were confined within a specific cell cycle phase. For example, cells in M-phase separated away from the other cell cycle phases along the UMAP1 axis **(Supp Fig. 2D)**. Second, other DDRs were defined by a combination of cell cycle phase and treatment. For example, cells in G1 are separated into two clusters. One cluster was mostly occupied by UVC-treated cells while the other was a mixture of IR and UVC-treated with untreated cells **(Supp. Fig. 2D, 2E)**. Third, cells exposed to the same treatment have distinct DDRs in the same cell cycle phase. For example, cells with etoposide resided in two distinct S-phase clusters. These data show that measuring and integrating DDR proteins with cell cycle at single cell resolution can reveal subpopulations with significant and previously unrecognized differences between perturbations.

**Measuring carboplatin-mediated DDR in HeLa cells**

To ask whether perturbation with carboplatin also resulted in distinct cell-specific responses, we treated HeLa cells with carboplatin for 24h and 48h. As previously reported, HeLa cells were highly resistant to carboplatin ($IC_{50}$ 317 μM) **(Supp. Fig. 3C)** [17]. UMAP embedding revealed a clear DDR, primarily in S-phase cells, to carboplatin treatment **(Supp. Fig. 3D-G)**. These data confirmed that carboplatin, like IR and UVC, can induce distinct cell-specific responses in asynchronously growing cells.

Supplementary Figures

### Coordinated and conserved DNA damage protein modules define responses to carboplatin in ovarian cancer cell lines

Jacob S. Bedia, Ying-Wen Huang, Antonio Delgado Gonzalez, Veronica D. Gonzalez, Ionut-Gabriel Funingana, Zainab Rahil, Alyssa Mike, Alexis Lowber, Maria Vias, Alan Ashworth[#], James D. Brenton[#], Wendy J. Fantl[#,*]

### co-senior authors

#### Contents

Supplementary Figures 1-9

Supplementary Tables 1-4

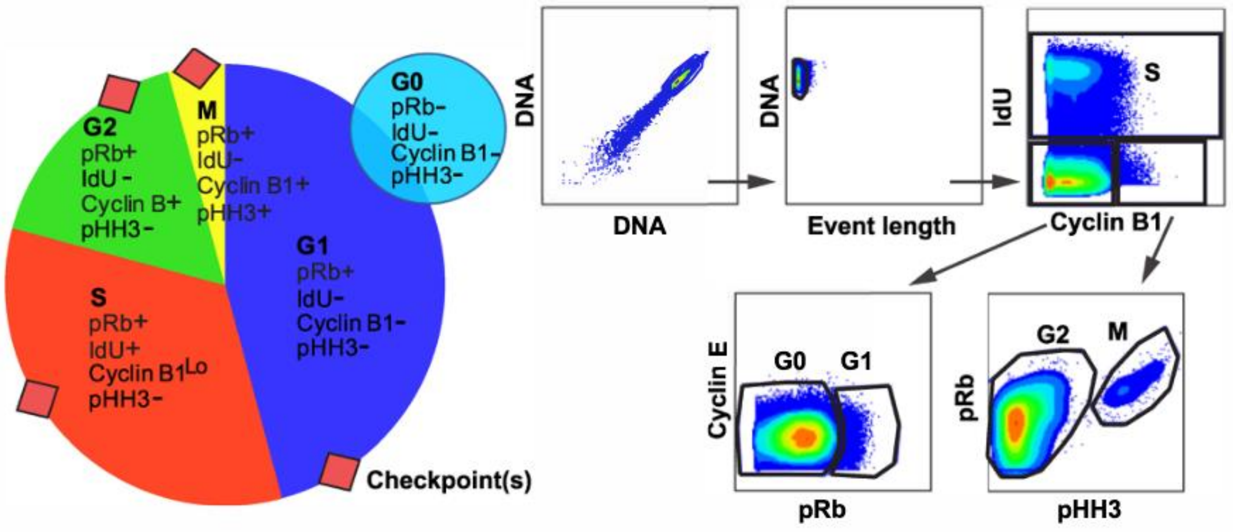


**Supp. Fig. 1. Gating scheme for cell cycle phases.** Pie-chart shows protein expression profiles by cell cycle phase. Gating scheme shows strategy for identifying cell cycle phases from levels of IdU, Cyclin B1, pRb, pHH3, and Cyclin E.

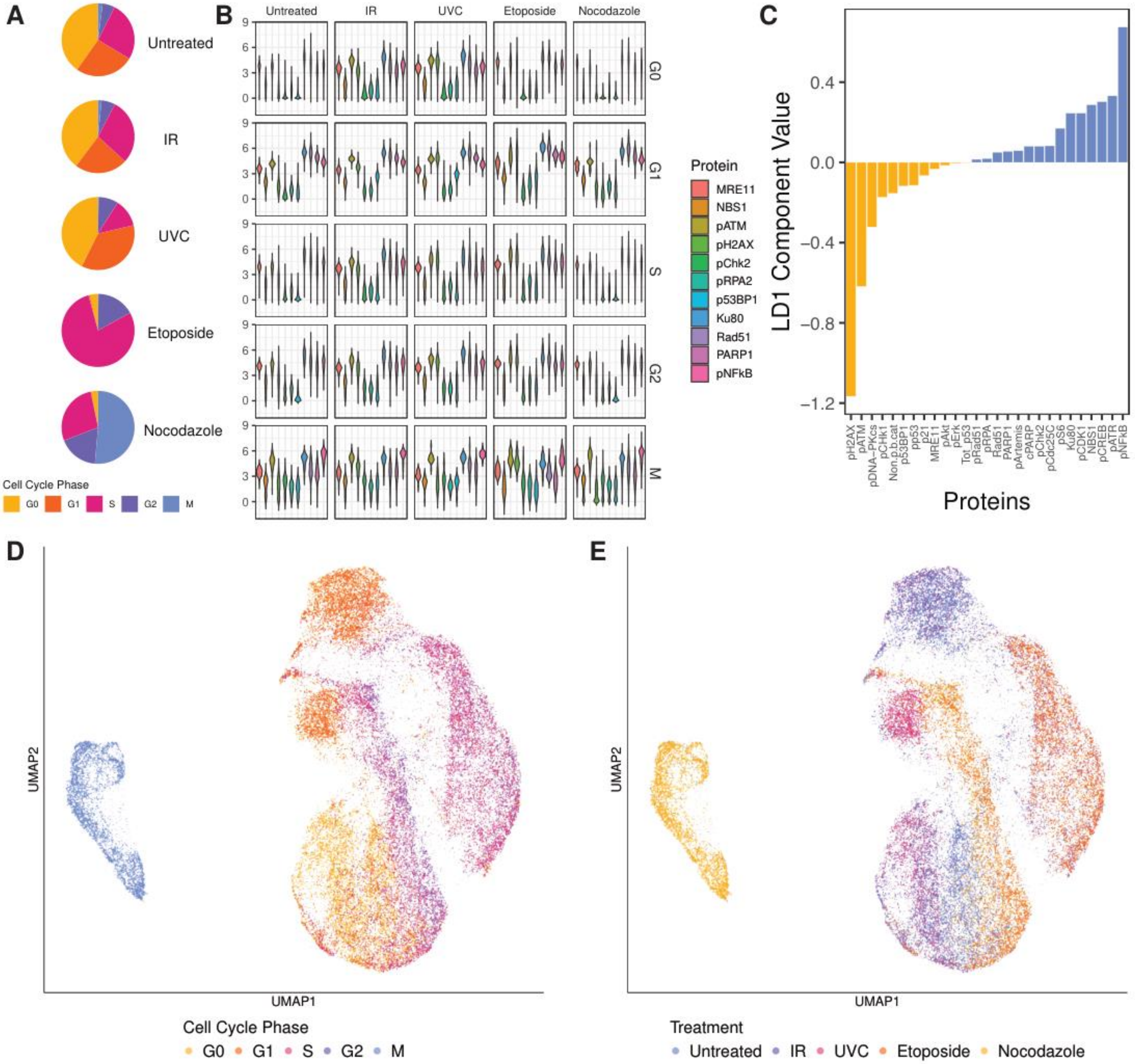

A
Untreated
IR
UVC
Etoposide
Nocodazole
Cell Cycle Phase
G0
G1
S
G2
M
B
Untreated
IR
UVC
Etoposide
Nocodazole
G0
G1
S
G2
M
Protein
MRE11
NBS1
pATM
pH2AX
pChk2
pRPA2
p53BP1
Ku80
Rad51
PARP1
pNFkB
C
LD1 Component Value
0.4
0.0
−0.4
−0.8
−1.2
pH2AX
pATM
pDNA−PKcs
pCHk1
Non.p.b.cat
p53BP1
pp53
p21
MRE11
pAkt
pErk
Tot_p53
pRad51
pRPA
Rad51
PARP1
pArtemis
cPARP
pChk2
pCdc25C
pS6
Ku80
pCDK1
NBS1
pCREB
pATR
pNFkB
Proteins
D
UMAP2
UMAP1
Cell Cycle Phase
G0
G1
S
G2
M
E
UMAP2
UMAP1
Treatment
Untreated
IR
UVC
Etoposide
Nocodazole

**Supp. Fig. 2. Validation of DDR CyTOF antibody panel using HeLa cells.** Cells were untreated (UT) or exposed to ionizing radiation (IR), ultraviolet C (UVC), nocodazole or etoposide. **A.** Pie charts depict cell frequency distributions across cell cycle phases. **B.** Violin plots show expression patterns of selected DDR proteins in cells across each cell cycle phase for each treatment. **C.** First linear discriminant function loadings show proteins with the most discriminating power between the five conditions. **D.** UMAP embedding of single cell HeLa data colored by cell cycle phase shows cells in same cell cycle phase with distinct DDRs. **E.** UMAP embedding of single cell HeLa data colored by treatment shows cells undergoing same treatment with distinct DDRs.

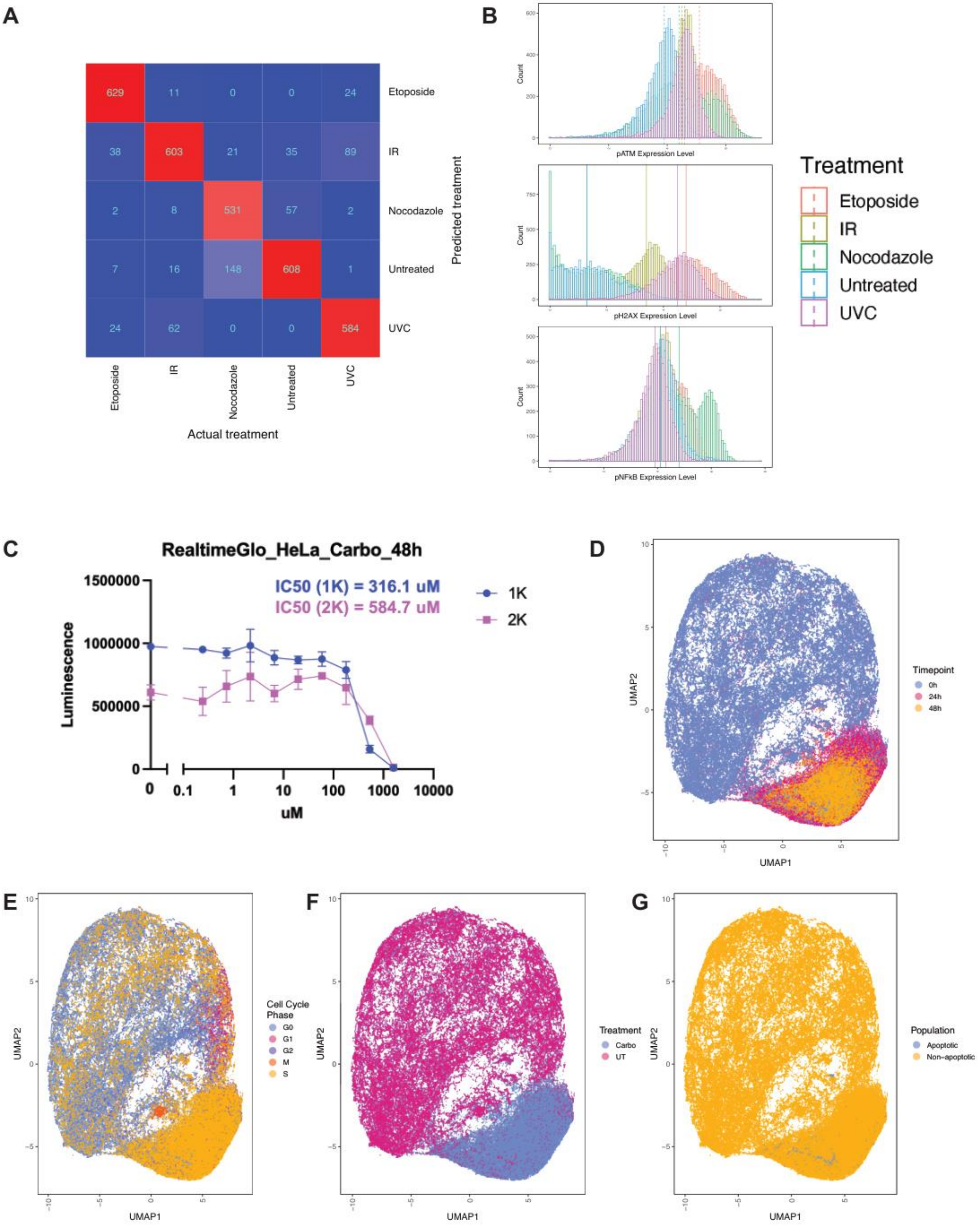
A
629
11
0
0
24
Etoposide
38
603
21
35
89
IR
2
8
531
57
2
Nocodazole
7
16
148
608
1
Untreated
24
62
0
0
584
UVC
Predicted treatment
Etoposide
IR
Nocodazole
Untreated
UVC
Actual treatment
B
Count
pATM Expression Level
Count
pH2AX Expression Level
Count
pNFkB Expression Level
Treatment
Etoposide
IR
Nocodazole
Untreated
UVC
C
RealtimeGlo_HeLa_Carbo_48h
IC50 (1K) = 316.1 uM
IC50 (2K) = 584.7 uM
1K
2K
Luminescence
1500000
1000000
500000
0
0
0.1
1
10
100
1000
10000
uM
D
UMAP2
UMAP1
Timepoint
0h
24h
48h
E
UMAP2
UMAP1
Cell Cycle Phase
G0
G1
G2
M
S
F
UMAP2
UMAP1
Treatment
Carbo
UT
G
UMAP2
UMAP1
Population
Apoptotic
Non-apoptotic

**Supp. Fig. 3. DNA damage responses of HeLa cells to diverse treatments. A**. Confusion matrix shows classification accuracy of linear discriminant functions based on DDR protein expression levels in a test set of 7000 HeLa cells. **B.** Histograms show differential expression of pATM, pH2AX, and pNFkB across the five treatments. Vertical lines denote median protein level for cells subjected to each treatment. **C.** Dose-response curves for HeLa cells treated with carboplatin, with IC50 by seeding density shown. Colors denote initial seeding density. **D-F.** UMAP of HeLa cells subjected to a carboplatin treatment time course show diverse DDRs to treatment. UMAPs are colored by timepoint, cell cycle phase, treatment, or cell fate, respectively.

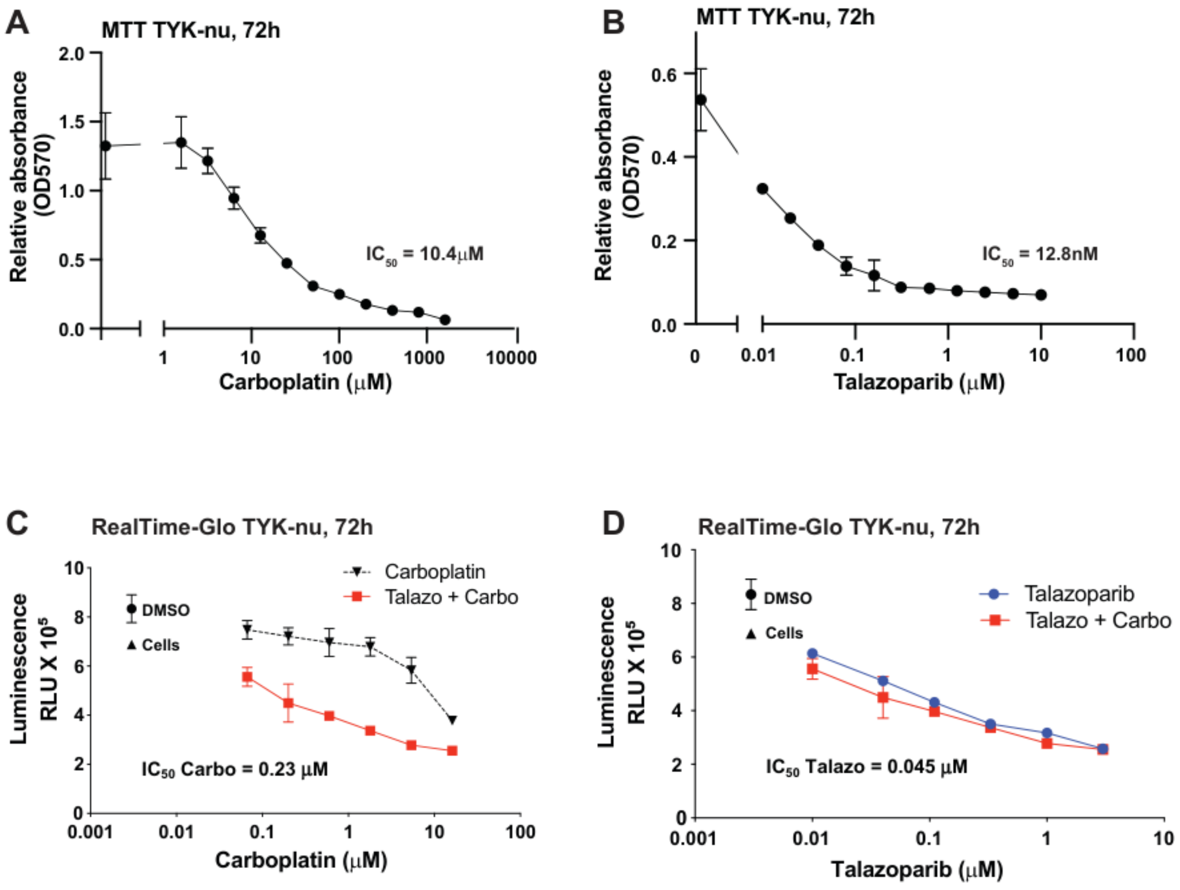


**Supp. Fig. 4. Dose response curves for TYK-nu cells treated with carboplatin and talazoparib.** Dose response curves for TYK-nu cells treated with carboplatin and talazoparib. **A.** MTT assay for carboplatin alone. **B.** MTT assay for talazoparib alone. **C.** Real time Glo assay for drug combination scaled for carboplatin. **D.** Real time Glo assay for drug combination scaled for talazoparib.

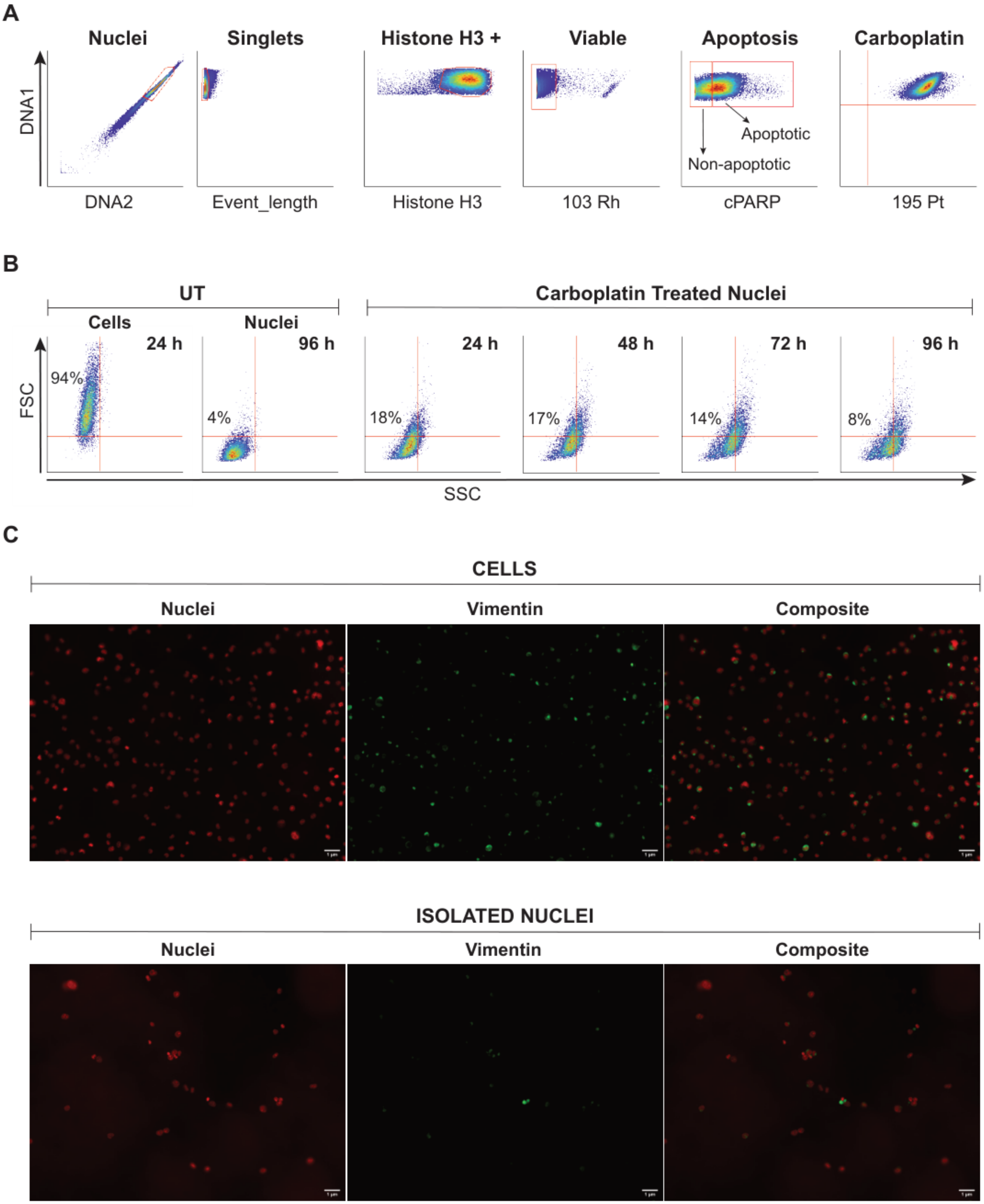
A
Nuclei
Singlets
Histone H3 +
Viable
Apoptosis
Carboplatin
DNA1
DNA2
Event_length
Histone H3
103 Rh
Apoptotic
Non-apoptotic
cPARP
195 Pt
B
UT
Carboplatin Treated Nuclei
Cells
Nuclei
24 h
96 h
24 h
48 h
72 h
96 h
FSC
94%
4%
18%
17%
14%
8%
SSC
C
CELLS
Nuclei
Vimentin
Composite
ISOLATED NUCLEI
Nuclei
Vimentin
Composite

**Supp. Fig. 5. Validation of single intact nuclei isolation protocol. A.** Gating scheme for identification of apoptotic and non-apoptotic populations in single, intact nuclei. **B.** Biaxial plots from flow cytometry experiments with TYK-nu cells and nuclei. Untreated nuclei show a more uniform forward scatter with carboplatin treated nuclei showing a similar forward scatter pattern. **C.** Immunofluorescence of TYK-nu cells and nuclei show significantly decreased vimentin expression in isolated nuclei.

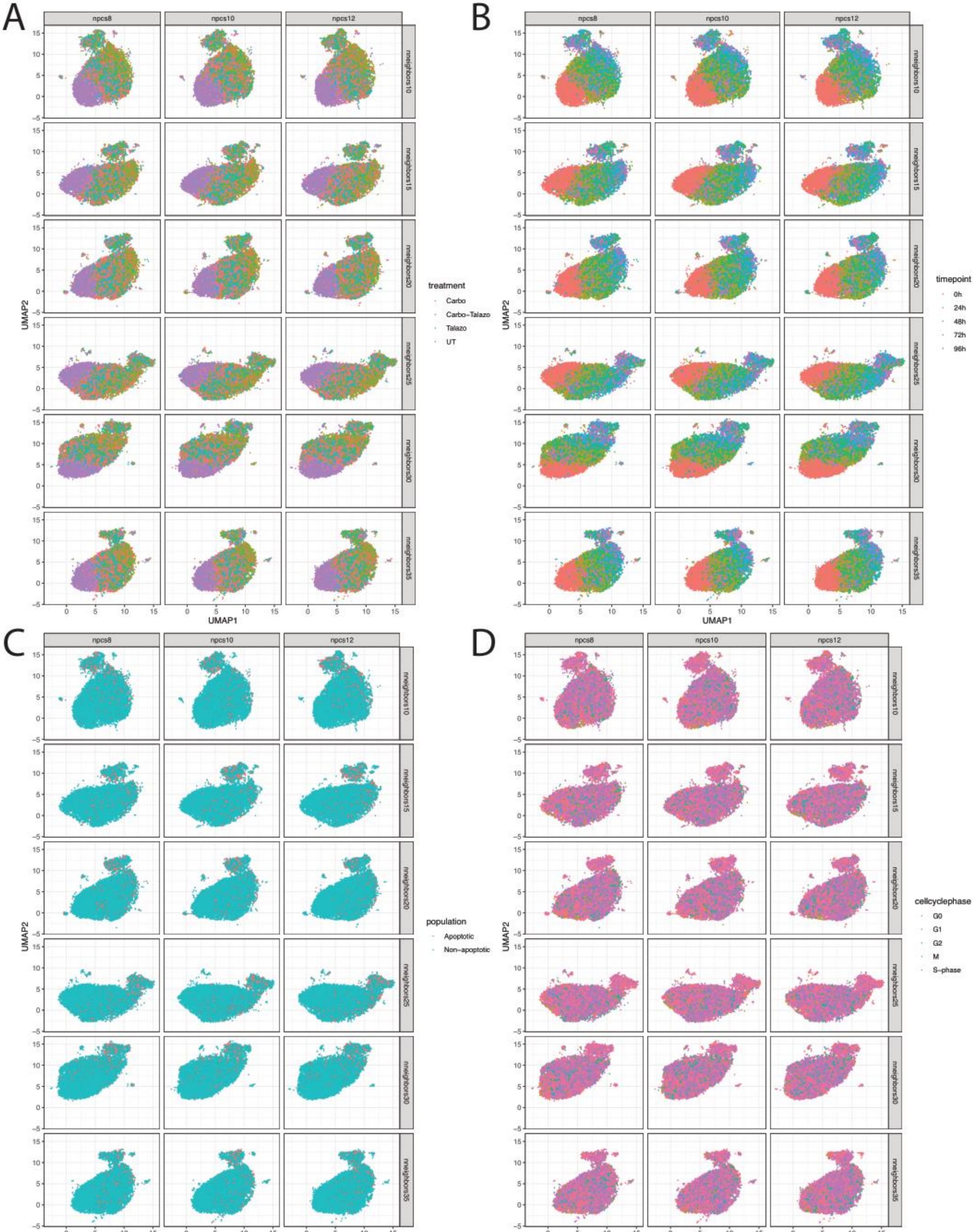

A
B
C
D
npcs8
npcs10
npcs12
nneighbors10
nneighbors15
nneighbors20
nneighbors25
nneighbors30
nneighbors35
UMAP1
UMAP2
treatment
Carbo
Carbo-Talazo
Talazo
UT
timepoint
0h
24h
48h
72h
96h
population
Apoptotic
Non-apoptotic
cellcyclephase
G0
G1
G2
M
S-phase

**Supp. Fig. 6. Stability of dimensionality reduction parameters for TYK-nu cells.** Parameter search for number of principal components and number of neighbors in k-nearest neighbor graph with changes in macroscopic structure of single cell data visualized by UMAP for **A.** Treatment. **B.** Timepoint. **C.** Population. **D.** Cell cycle phase.

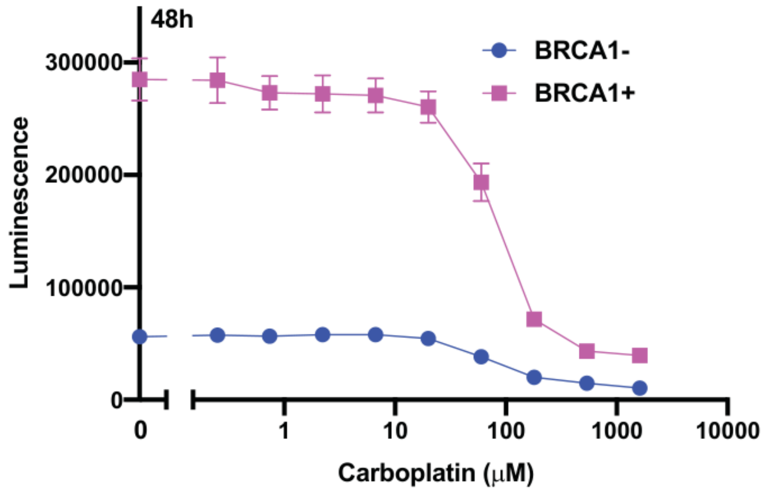


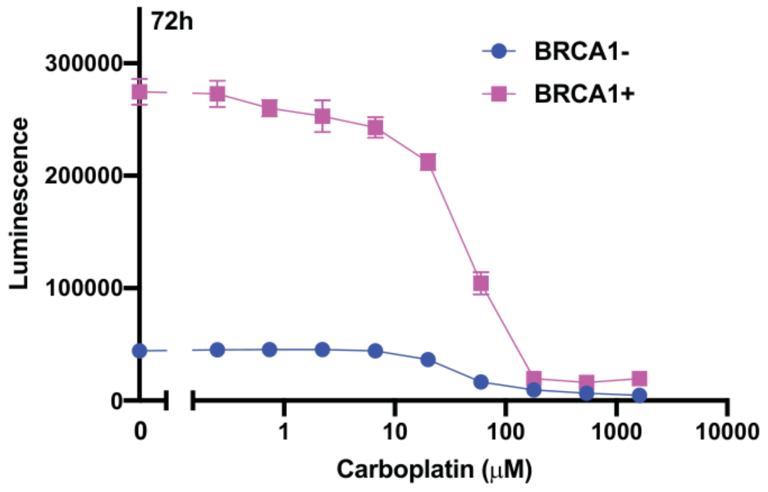


**Supp. Fig. 7. Dose response curves for UWB cells treated with carboplatin.**

Real time Glo assay for UWB cell lines treated with carboplatin for 48 or 72 hr.

Timepoint denotes duration of treatment

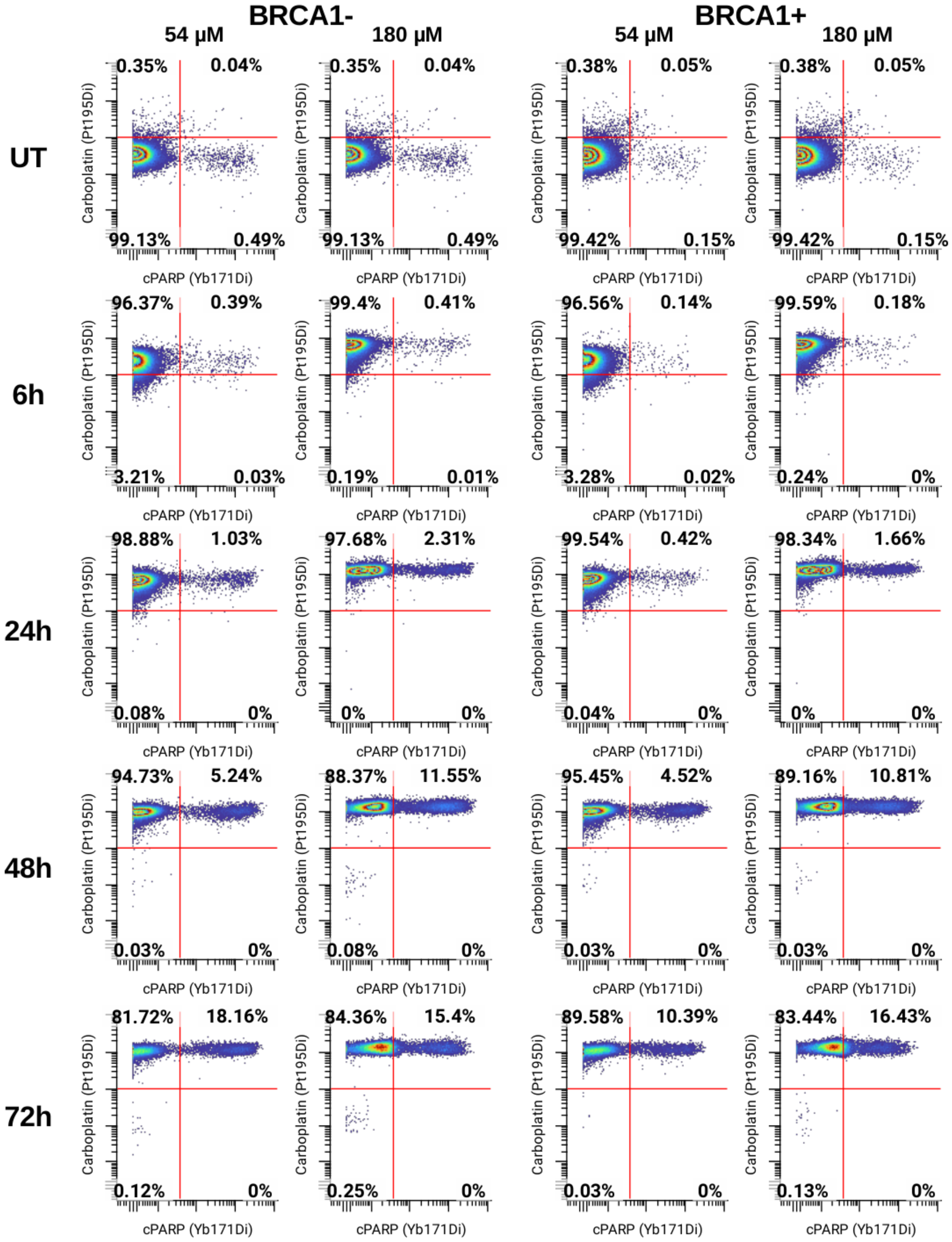
BRCA1-
BRCA1+
54 µM
180 µM
54 µM
180 µM
UT
6h
24h
48h
72h
Carboplatin (Pt195Di)
cPARP (Yb171Di)
UT, BRCA1- 54 µM: 0.35% 0.04% 99.13% 0.49%
UT, BRCA1- 180 µM: 0.35% 0.04% 99.13% 0.49%
UT, BRCA1+ 54 µM: 0.38% 0.05% 99.42% 0.15%
UT, BRCA1+ 180 µM: 0.38% 0.05% 99.42% 0.15%
6h, BRCA1- 54 µM: 96.37% 0.39% 3.21% 0.03%
6h, BRCA1- 180 µM: 99.4% 0.41% 0.19% 0.01%
6h, BRCA1+ 54 µM: 96.56% 0.14% 3.28% 0.02%
6h, BRCA1+ 180 µM: 99.59% 0.18% 0.24% 0%
24h, BRCA1- 54 µM: 98.88% 1.03% 0.08% 0%
24h, BRCA1- 180 µM: 97.68% 2.31% 0% 0%
24h, BRCA1+ 54 µM: 99.54% 0.42% 0.04% 0%
24h, BRCA1+ 180 µM: 98.34% 1.66% 0% 0%
48h, BRCA1- 54 µM: 94.73% 5.24% 0.03% 0%
48h, BRCA1- 180 µM: 88.37% 11.55% 0.08% 0%
48h, BRCA1+ 54 µM: 95.45% 4.52% 0.03% 0%
48h, BRCA1+ 180 µM: 89.16% 10.81% 0.03% 0%
72h, BRCA1- 54 µM: 81.72% 18.16% 0.12% 0%
72h, BRCA1- 180 µM: 84.36% 15.4% 0.25% 0%
72h, BRCA1+ 54 µM: 89.58% 10.39% 0.03% 0%
72h, BRCA1+ 180 µM: 83.44% 16.43% 0.13% 0%

**Supp. Fig. 8. UWB cell fate responses to treatment at low and high concentrations of carboplatin.** Biaxial plots of intracellular platinum levels (Pt-195) vs. cPARP in live (Rh103-) single UWB cells treated with carboplatin in 72 hr time course.

Percentages denote the percentage of live cells in each quadrant of quad gate.

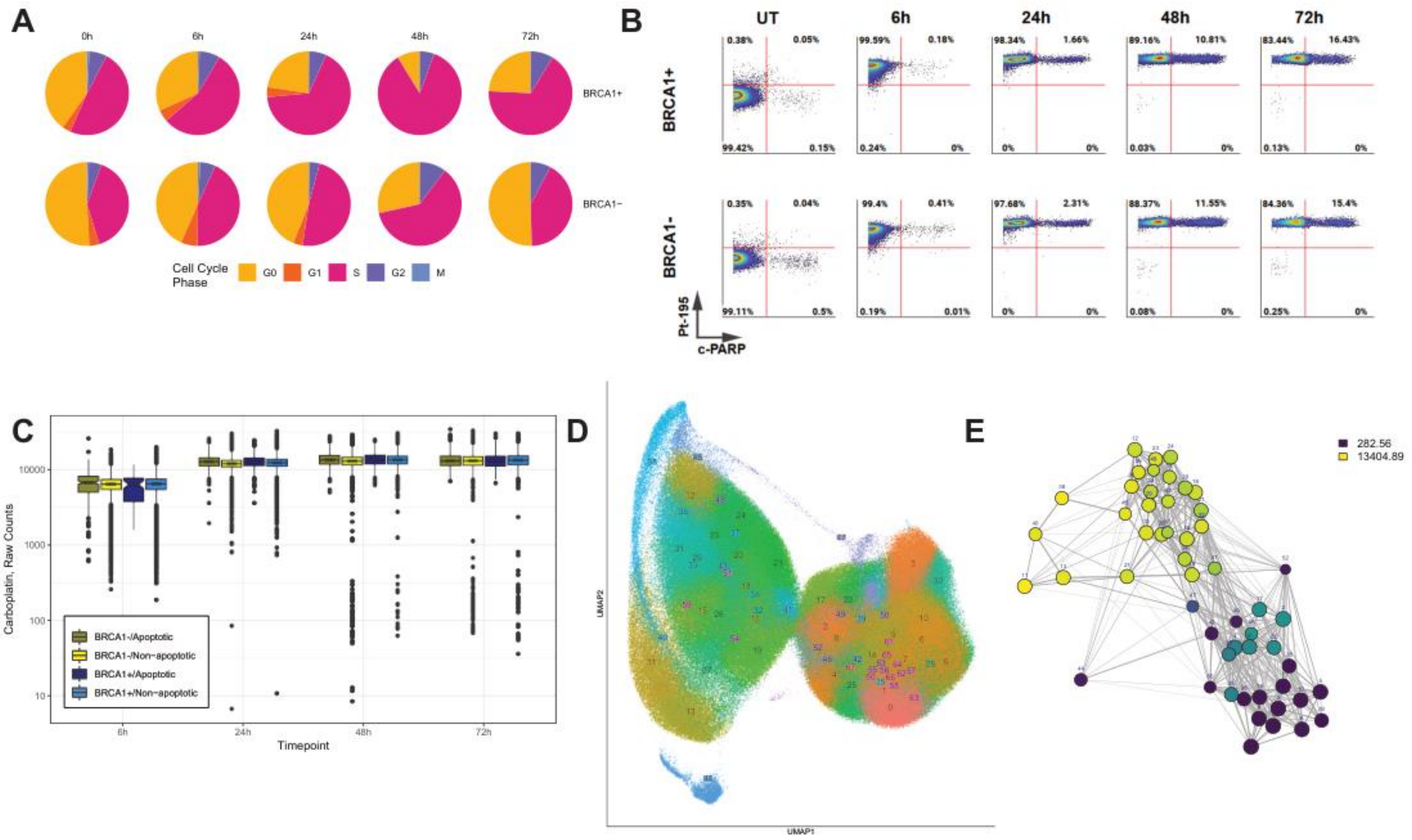


**Supp. Fig. 9. UWB single cell analysis. A.** Pie charts depict cell frequency distributions across cell cycle phases. **B.** Biaxial plots of Pt-195 vs. c-PARP show proportion of apoptotic cells in response to treatment over time. **C.** Box and whisker plot showing platinum uptake in single intact cells. Boxes are colored by cell fate (apoptotic or non-apoptotic) or cell line. **D.** UMAP of UWB single cell data colored by cluster assignment. **E.** PAGA graph colored by median platinum levels in each cluster.

#### Supplementary Table 1

| Isotope | Antigen | Clone | Phosphorylation site | Vendor | Positive control cell line | Negative control cell line |
|---|---|---|---|---|---|---|
| Y89 | Cyclin A | BF683 | | BD | HCT116 - Nocodazole 18h | HCT116 - DMSO 0.005% |
| In113 | Cyclin E | HE12 | | Life Technology | HCT116 - Hydroxyurea 20h | HCT116 - H2O |
| In115 | Vimentin | D21H3 | | CST | NIH3T3 | OVCAR3 |
| Pr141 | pRPA32/RPA2 | Polyclonal | T21 | Abcam | HELA - Etoposide 24h | HELA - DMSO (0.05%) |
| Nd142 | pCDK1/cdc2 | 10A11 | Y15 | CST | HCT116 - Nocodazole 18h | HCT116 - DMSO (0.005%) |
| Nd143 | pCHK2 | C13C1 | T68 | CST | HCT116 - Nocodazole 18h | HCT116 - DMSO (0.005%) |
| Nd145 | NBS1 | 34/NBS1 | | BD | K562 | PBMC- CD3+ T cells |
| Nd146 | pATM | 10H11.E12 | S1981 | Millipore | HELA - Etoposide 24h | HELA - DMSO (0.05%) |
| Sm147 | pH2AX | JBW301 | S139 | Millipore | HELA - Etoposide 24h | HELA - DMSO (0.05%) |
| Nd148 | Cyclin B1 (total) | GNS-1 | | BD | HCT116 - Nocodazole 18h | HCT116 - DMSO 0.005% |
| Sm149 | pNFkB | K10-895.12.50 | S529 | BD | Jurkat- TNFa 15min | Jurkat- PBS |
| Nd150 | pAurora A/B/C | D13A11 | | CST | HCT116 - Nocodazole 18h | HCT116 - DMSO (0.005%) |
| Eu151 | pERK | D13.14.4E | pT202/ pY204 | CST | Jurkat- Pervanadate 15min | Jurkat - DMSO (0.0625%) |
| Sm152 | Ku80 | 7/Ku80 | | BD | HELA | NIH 3T3 |
| Eu153 | p-p53 | 16G8 | S15 | CST | HCTI 16 - Hydroxyurea 20h | HCT116 - H20 |
| Sm154 | pCdc25C | E190 | S216 | Abcam | HCT116 - Nocodazole 18h S phase | HCT116 - Nocodazole 18h M phase |
| Gd155 | Ki67 (total) | B56 | | BD | OVCAR4 | OVKATE |
| Gd156 | pArtemis | Polyclonal | S516 | Abcam | HELA | PBMC- CD3+ T cells |
| Gd157 | pRad51 | Polyclonal | Y315 | Sigma Aldrich | TykNu | PBMC- CD3+ T cells |
| Gd158 | E-Cadherin | 67A4 | | Biolegend | OVCAR3 | U937 |
| Tb159 | pAkt | D9E | S473 | CST | Kuramochi, serum starved O/N, EGF 10min | Kuramochi - serum starved O/N, PBS |
| Gd160 | p53BP1 | Polyclonal | S1778 | CST | HCT116 - Nocodazole 18h S phase | HCT116 - Nocodazole 18h M phase |
| Dy161 | c-Myc | D84C12 | | CST | TykNu | MCF7 |
| Dy162 | p21 | SXM30 | | BD | MCF7 | HELA |
| Dy164 | PCNA | PC10 | | BD | Jurkat | PBMC- CD14- Leukocytes |
| Ho165 | pRb | J112-906 | S807/ S811 | BD | OVSAHO | PBMC- CD3+ T cells |
| Er166 | pATR | Polyclonal | T1989 | GeneTEx | HCT116 - Nocodazole 18h | HCT116 - DMSO (0.005%) |
| Er167 | Rad51 | Polyclonal | | EMD Millipore | HELA | PBMC- CD3+ T cells |
| Er168 | PARP1 | 839120 | | R&D systems | NALM6 | PBMC- CD14+ Monocytes |
| Tm169 | pCHK1 | 133D3 | S345 | CST | HELA - Hydroxyurea 1h pH2AX+ | HELA - Hydroxyurea 1h pH2AX- |
| Er170 | Non-phospho beta catenin | D13A1 | | CST | Kuramochi | U937 |
| Yb171 | cPARP | F21-852 | cleaved N214 | BD | U937 - Etoposide 24hr | U937 - DMSO (0.1%) |
| Yb172 | prpS6 | N7-548 | S235/ S236 | BD | OVCAR4 | U937 |
| Yb173 | pDNA-PKcs | 10B1 | T2609 | Biolegend | HELA-UVC 100J/m2 | HELA - Untreated |
| Lu175 | Total p53 | 1C12 | | CST | Kuramochi | U937 |
| Yb176 | pHH3 | HTA28 | S28 | Biolegend | HCT116 - Nocodazole 18h | HCT116 - DMSO (0.005%) |
| Bi209 | MRE11 | 18/MRE11 | | BD | K562 | PBMC- CD14- Leukocytes |

**Supp. Table 1. CyTOF validated antibody panel.** Antibody clones were selected based on their performance in fluorescence-based flow cytometry. All metal conjugations were carried out in- house. Working concentrations were optimized by titration against positive and negative controls (right-hand columns).

#### Supplementary Table 2

| Cell Line | Gene | Chr | Start Position | End Position | Variant Classification | Variant Annotation | Variant Type | Protein Change |
|---|---|---|---|---|---|---|---|---|
| TYKnu | ACKR3 | 2 | 237489798 | 237489798 | Silent | silent | SNP | p.V230V |
| TYKnu | ACKR3 | 2 | 237489924 | 237489924 | Silent | silent | SNP | p.V272V |
| TYKnu | ARID1A | 1 | 27023164 | 27023174 | Frame_Shift_Del | damaging | DEL | p.GGGP91fs |
| TYKnu | ATRX | X | 76776298 | 76776298 | Missense_Mutation | other non-conserving | SNP | p.I2390V |
| TYKnu | CHD1L | 1 | 146756029 | 146756029 | Missense_Mutation | other non-conserving | SNP | p.H571Y |
| TYKnu | DOT1L | 19 | 2226473 | 2226473 | Missense_Mutation | other non-conserving | SNP | p.F1318C |
| TYKnu | DYRK3 | 1 | 206821588 | 206821588 | Missense_Mutation | other non-conserving | SNP | p.S349G |
| TYKnu | ERCC4 | 16 | 14038684 | 14038684 | Missense_Mutation | other non-conserving | SNP | p.R670Q |
| TYKnu | FXR1 | 3 | 180630510 | 180630510 | Frame_Shift_Del | damaging | DEL | p.G13fs |
| TYKnu | GCNA | X | 70823819 | 70823819 | Missense_Mutation | other non-conserving | SNP | p.D231G |
| TYKnu | HDAC10 | 22 | 50689654 | 50689654 | De_novo_Start_OutOfFrame | damaging | SNP | NA |
| TYKnu | HELQ | 4 | 84376682 | 84376682 | Silent | silent | SNP | p.T55T |
| TYKnu | IRF3 | 19 | 50165278 | 50165278 | Silent | silent | SNP | p.S303S |
| TYKnu | MSH3 | 5 | 80071511 | 80071511 | Splice_Site | damaging | SNP | NA |
| TYKnu | MSH6 | 2 | 48033981 | 48033982 | Frame_Shift_Ins | damaging | INS | p.-1357fs |
| TYKnu | NPM1 | 5 | 170827189 | 170827189 | Missense_Mutation | other non-conserving | SNP | p.A186V |
| TYKnu | PLA2R1 | 2 | 160843823 | 160843823 | Silent | silent | SNP | p.G627G |
| TYKnu | RAD51AP2 | 2 | 17699222 | 17699222 | Missense_Mutation | other non-conserving | SNP | p.G154E |
| TYKnu | SETD1A | 16 | 30972611 | 30972611 | Missense_Mutation | other non-conserving | SNP | p.Q90H |
| TYKnu | SETD1A | 16 | 30976605 | 30976605 | Silent | silent | SNP | p.E514E |
| TYKnu | TAOK2 | 16 | 29998131 | 29998131 | Missense_Mutation | other non-conserving | SNP | p.I846M |
| TYKnu | TELO2 | 16 | 1550423 | 1550423 | Missense_Mutation | other non-conserving | SNP | p.V360I |
| TYKnu | TICRR | 15 | 90166966 | 90166966 | Missense_Mutation | other non-conserving | SNP | p.P1142L |
| TYKnu | TP53 | 17 | 7578406 | 7578406 | Missense_Mutation | other non-conserving | SNP | p.R175H |
| TYKnu | VAV3 | 1 | 108160227 | 108160227 | Nonsense_Mutation | damaging | SNP | p.G648* |
| UWB1289 | ABL1 | 9 | 133730285 | 133730285 | Silent | silent | SNP | p.T117T |
| UWB1289 | BAZ1B | 7 | 72892617 | 72892617 | Missense_Mutation | other non-conserving | SNP | p.K392E |
| UWB1289 | BRCA1 | 17 | 41245073 | 41245073 | Frame_Shift_Del | damaging | DEL | p.D825fs |
| UWB1289 | DYRK1B | 19 | 40319150 | 40319150 | Silent | silent | SNP | p.Y198Y |
| UWB1289 | E2F7 | 12 | 77449707 | 77449707 | Missense_Mutation | other non-conserving | SNP | p.R99S |
| UWB1289 | PARG | 10 | 51050038 | 51050038 | Silent | silent | SNP | p.L348L |
| UWB1289 | SETD1A | 16 | 30991502 | 30991502 | Missense_Mutation | other non-conserving | SNP | p.W1465C |
| UWB1289 | TP53 | 17 | 7578222 | 7578223 | Frame_Shift_Del | damaging | DEL | p.R209fs |
| UWB1289 | XAB2 | 19 | 7688729 | 7688729 | Missense_Mutation | other non-conserving | SNP | p.Q336R |

**Supp. Table 2. Mutations in DDR genes across cell lines.** Mutation profiles obtained from DepMap for cell lines used in this study (TYK-nu, UWB1289) . See Methods for details of table generation.

**Supplementary Table 3**

| Treatment | Population | Timepoint | Cell Cycle Phase | | | | |
|---|---|---|---|---|---|---|---|
| | | | G0 | G1 | S-phase | G2 | M |
| Carbo | Apoptotic | 24h | 207 | 5 | 63 | 37 | 39 |
| Carbo | Apoptotic | 48h | 709 | 5 | 494 | 105 | 28 |
| Carbo | Apoptotic | 72h | 2211 | 2 | 486 | 90 | 13 |
| Carbo | Apoptotic | 96h | 2967 | 2 | 426 | 165 | 14 |
| Carbo | Non-apoptotic | 24h | 14202 | 2699 | 52361 | 6842 | 2000 |
| Carbo | Non-apoptotic | 48h | 7933 | 1343 | 67402 | 5884 | 711 |
| Carbo | Non-apoptotic | 72h | 5091 | 161 | 27340 | 4043 | 160 |
| Carbo | Non-apoptotic | 96h | 4569 | 186 | 13747 | 4332 | 95 |
| Carbo-Talazo | Apoptotic | 24h | 191 | 3 | 191 | 89 | 19 |
| Carbo-Talazo | Apoptotic | 48h | 4090 | 7 | 1987 | 1772 | 23 |
| Carbo-Talazo | Apoptotic | 72h | 6808 | 4 | 1205 | 514 | 10 |
| Carbo-Talazo | Apoptotic | 96h | 9599 | 5 | 1123 | 760 | 36 |
| Carbo-Talazo | Non-apoptotic | 24h | 5031 | 895 | 32204 | 1700 | 308 |
| Carbo-Talazo | Non-apoptotic | 48h | 6383 | 324 | 37099 | 4247 | 82 |
| Carbo-Talazo | Non-apoptotic | 72h | 6042 | 191 | 13333 | 4723 | 41 |
| Carbo-Talazo | Non-apoptotic | 96h | 4978 | 89 | 7254 | 5611 | 33 |
| Talazo | Apoptotic | 24h | 283 | 11 | 267 | 168 | 41 |
| Talazo | Apoptotic | 48h | 2884 | 10 | 1416 | 1185 | 14 |
| Talazo | Apoptotic | 72h | 1373 | 0 | 216 | 118 | 6 |
| Talazo | Apoptotic | 96h | 4379 | 15 | 647 | 133 | 9 |
| Talazo | Non-apoptotic | 24h | 8626 | 1725 | 53066 | 3620 | 631 |
| Talazo | Non-apoptotic | 48h | 7600 | 782 | 37841 | 4184 | 307 |
| Talazo | Non-apoptotic | 72h | 1688 | 57 | 4161 | 829 | 36 |
| Talazo | Non-apoptotic | 96h | 7913 | 541 | 19818 | 6049 | 410 |
| UT | Apoptotic | 0h | 563 | 3 | 63 | 30 | 20 |
| UT | Non-apoptotic | 0h | 54655 | 2035 | 77708 | 13975 | 7687 |

**Supp. Table 3.** TYK-nu cell counts by cell cycle phase, timepoint, cell fate, and treatment.

**Supplementary Table 4**

| Treatment | Population | Timepoint | Cell Cycle Phase | | | | |
|---|---|---|---|---|---|---|---|
| | | | G0 | G1 | S-phase | G2 | M |
| Carbo | Apoptotic | 24h | 675 | 16 | 1869 | 169 | 3 |
| Carbo | Apoptotic | 48h | 787 | 0 | 9201 | 212 | 2 |
| Carbo | Apoptotic | 6h | 179 | 14 | 84 | 36 | 5 |
| Carbo | Apoptotic | 72h | 381 | 1 | 3302 | 63 | 0 |
| Carbo | Non-apoptotic | 24h | 50807 | 5875 | 89959 | 8397 | 248 |
| Carbo | Non-apoptotic | 48h | 16328 | 208 | 64836 | 7076 | 37 |
| Carbo | Non-apoptotic | 6h | 49583 | 7183 | 65424 | 9184 | 987 |
| Carbo | Non-apoptotic | 72h | 12861 | 21 | 18840 | 2897 | 11 |
| UT | Apoptotic | 0h | 497 | 11 | 72 | 27 | 4 |
| UT | Non-apoptotic | 0h | 99308 | 7842 | 100253 | 13288 | 1782 |

**Supp. Table 4.** UWB cell counts by cell cycle phase, timepoint, cell fate, and treatment.
